# Comparing the signaling and transcriptome profiling landscapes of human iPSC-derived and primary rat neonatal cardiomyocytes

**DOI:** 10.1101/2022.09.06.506781

**Authors:** Kyla Bourque, Jace Jones-Tabah, Darlaine Pétrin, Ryan D. Martin, Terence E. Hébert

**Author notes:** Address correspondence to: Dr. Terence E. Hébert, PhD, Department of Pharmacology and Therapeutics, McGill University, 3655 Promenade Sir-William-Osler, Room 1325, Montréal, Québec, H3G 1Y6, Canada.

## Abstract

The inaccessibility of human cardiomyocytes significantly hindered years of cardiovascular research efforts. Post-mortem tissue or biopsies from diseased patients, which remain scarcely available, rendered it possible to study end-stage heart disease yet the inclusion of healthy human cardiac materials for basic science research was beyond reach. To overcome these limitations, non-human cell sources were used as proxies to study heart function and associated diseases. Rodent models became increasingly acceptable surrogates to model the human heart either *in vivo* or through *in vitro* cultures. More recently, due to concerns regarding animal to human translation, including cross-species differences, the use of human inducible stem cell derived cardiomyocytes presented a renewed opportunity. We think it necessary to conduct a comparative study, assessing cellular signalling through cardiac G protein-coupled receptors and bulk transcriptomics of traditional rat neonatal cardiomyocytes and human iPSC-CMs. Genetically-encoded biosensors were used to interrogate nuclear protein kinase A (PKA) and extracellular signal-regulated kinase 1/ 2 (ERK_1/2_) in rat and human-derived cardiomyocyte populations. To increase data granularity, a single-cell analytical approach was conducted for an in-depth examination of existing differences between both *in vitro* cardiomyocyte models. Using automated high content microscopy, our analyses of nuclear PKA and ERK_1/2_ signaling revealed distinct response clusters in rat and human CMs. In line with this, bulk RNA-seq demonstrated key differences regarding the expression patterns of GPCRs, G proteins and effectors. Overall, our study demonstrates that human stem cell derived models of the cardiomyocyte do provide significant advantages and should be taken advantage of.

## Introduction

Cardiac physiology is regulated, to some extent, by G protein-coupled receptors (GPCRs) including adrenergic, angiotensin and endothelin receptor systems. These receptors and their associated signalling effectors modulate cardiac contractility, force generation, vascular tone, as well as cardiomyocyte growth and survival [1, 2]. The activation of prototypical cardiac Gα_q_-coupled and Gα_s_-coupled GPCRs, α- and β-adrenergic receptors for instance, result in calcium mobilization or protein kinase A (PKA) activity, correspondingly influencing cardiomyocyte contractile responses [3]. Moreover, the activation of the extracellular signal-regulated kinase 1/2 (ERK_1/2_) downstream adrenergic and angiotensin input has relatedly been linked with cardiomyocyte viability and survival [4]. Yet, signals propagated by these kinases have also been associated with maladaptive cardiac remodelling and hypertrophic growth. The same receptors and effectors can thus drive adaptive and maladaptive cardiac functions. These complex signalling circuits regulating cardiomyocyte function and pathway activation have been extensively studied, in numerous model systems, as dysregulated activity downstream GPCR activation has been implicated in the progression of several heart muscle diseases.

Generally, the study of human cardiac physiology and pathophysiology has been challenging. The most notable barrier has been the inaccessibility of human cardiac tissue and cell sources for *in vitro* experimental studies. Human heart tissue obtained from diseased non-transplantable hearts only provide insight on end-stage heart disease and is not informative of disease evolution through time. The few adult primary human cardiomyocytes that may be rarely obtained, only last one or two days in culture, thereby limiting the types of experiments that can be conducted and reducing the chance to explore disease progression. Recently, a new isolation protocol has been described that was shown to maintain human primary cardiomyocytes in culture for 7 days as well as a procedure for successful cryopreservation and thawing [5]. Needless to say, the small quantities obtained still represent a substantial bottleneck and beyond the reach of most laboratories. To model the human heart, other *representative* cell sources were exploited, most prominently being cardiomyocyte cultures derived from rodents. With large litters and relatively low-cost for maintenance compared to large-animal models, the isolation of cardiomyocytes from these species provides millions of functional cells for study. Frequently, neonatal rat cardiomyocyte cultures are used as they can typically be kept in culture for 5-7 days compared to adult rodent cultures that tend to de-differentiate after 2-3 days *in vitro*. Numerous reports have demonstrated that cardiomyocytes isolated from mice and rat hearts are signalling competent and hypertrophy in response to known inducers [6, 7]. With certain genetic and phenotypic similarities to humans, rodents have helped researchers study numerous aspects of cardiac physiology. Nonetheless, rodent hearts do not completely recapitulate heart conduction and electrophysiological properties of the adult human heart. Most remarkably, while at rest, the human heart beats on average 60-70 beats per minute, while mice and rat hearts beat at 500-600 and 260-450 bpm, respectively [8]. Further, differences at the level of action potentials, myofilaments, isoforms of certain proteins and their corresponding phosphorylation status have also been reported [9]. Besides the species barrier, 5-7 days in culture represents a serious limitation when attempting to model long-term and slow onset diseases leading to heart failure. In addition, neonatal cultures may also limit the translatability of experimental results when modeling adult-onset diseases. This raises the ever-present underlying question, how well does the experimental data collected in neonatal rat cardiomyocytes translate to the *adult* human heart or at the level of the individual cardiomyocyte?

With the advent of human induced pluripotent stem cells (hiPSCs), the influence of crossspecies differences has been a topic of interest due to the establishment of robust protocols that permit the differentiation of hiPSCs towards mesodermal cell types like cardiomyocytes (CMs) (reviewed in [10, 11]). Human derived iPSC-CMs represent a clinically relevant cell population of high purity, quality and quantity [12]. These differentiated cells express the relevant cell- and tissue-specific receptors and ion channels compared to their *in vivo* counterparts. They can even be cryopreserved providing access to large numbers of cells with less batch-to batch variation [13]. Considering that certain lines of evidence suggest that cellular signalling networks that control myocardial function are cell- and tissue-type specific, it becomes vital to assess experimental endpoints in a human background. As such, we sought to examine whether there were differences between primary rodent and human derived cardiomyocytes through dual profiling of cellular signaling pathways and the transcriptome. Briefly, we measured pathway activation downstream of four cardiac relevant GPCR families, namely α_1_-, β_1_- and β_2_-adrenergic receptors (α_1_AR, β_1_- and β_2_AR), AT_1_R angiotensin, and ET_A_ endothelin receptors. We assessed signalling transduction pathways using genetically-encoded biosensors that measure nuclear PKA and ERK_1/2_ activity. Pathway activation was analyzed on a single-cell basis in order to parse out subtle differences within the overall rat neonatal or human iPSC-derived cardiomyocyte populations. To complement these assays, we also investigated gene expression via bulk transcriptomics using RNA-seq. Further, gene expression profiles of rat and iPSC-CMs were compared to HEK 293 cells as these have been a valuable resource to study the biological function of cardiac relevant GPCRs.

## Materials and Methods

### Reagents

Unless specified, all common laboratory reagents were purchased from Sigma-Aldrich. Drugs were purchased from various vendors as follows: forskolin (Bioshop, FRS393.5), phorbol-12-myristate-13-acetate (PMA-, Cedarlane 10008014-1), angiotensin II (Sigma, A9525), phenylephrine (Sigma Aldrich, P6126), endothelin-1 (Bachem, H6995.0001), isoproterenol (Sigma Aldrich, I6504), norepinephrine (Sigma Aldrich, A9512), epinephrine (Sigma Aldrich, E4375), ascorbic acid (Sigma Aldrich, A4544), acetic acid (Fisherbrand, 351270-212).

### Rat neonatal cardiomyocyte isolation and culture

All procedures involving animals were approved by the McGill University Animal Care Committee, in accordance with Canadian Council on Animal Care Guidelines. Sprague-Dawley dams with postnatal day 1-3 pups were purchased from Charles River, Saint-Constant QC, Canada. Complete litters, including both female and male neonatal pups were sacrificed by decapitation, as previously described [14–16]. Using forceps, whole hearts were isolated from the chest cavity and placed in cold HBSS (Wisent, 311-511-CL). Using surgical scissors, hearts were kept whole and cut 3-5 times to increase the surface area for overnight enzymatic digestion at 4 °C with 0.1% trypsin in HBSS. The next morning, the trypsin reaction was inhibited with 7% FBS supplemented DMEM low glucose + penicillin/streptomycin (P/S). Five serial collagenase digestions were then performed. Cell suspensions containing cardiomyocytes and cardiac fibroblasts from whole heart extracts were then seeded onto tissue-culture treated plastic 10-cm dishes. Non-cardiomyocyte cells such as fibroblasts can attach to plastic while cardiomyocytes do not adhere to plastic surfaces. The cell suspension after this incubation period is thus enriched for cardiomyocytes. After two 75-minute incubations, the cell suspension typically contains >90% cardiomyocytes and were seeded in black optical bottom 96-well plates (Thermo Scientific, 165305) coated with human plasma fibronectin + 0.1% gelatin in DMEM low glucose + 7% (vol/vol) FBS + P/S + 10 μM cytosine-β-d-arabinoside (AraC, Sigma Aldrich, C1768-500MG). The mitotic inhibitor, AraC, was added to prevent proliferation of remaining dividing fibroblasts. Cardiomyocytes were then maintained in a humidified atmosphere at 37 °C with 5% CO_2_. The next day, cardiomyocytes were washed three times with DMEM low glucose and exchanged with maintenance medium containing DMEM low glucose and universal ITS supplement composed of insulin, transferrin, and selenium (Wisent, 315-081-QL) prior to being transduced with adeno-associated viruses carrying the genetically-encoded biosensor of interest. For the RNAseq experiment, RNCMs were seeded in a 6-well dish coated with human plasma fibronectin + 0.1% gelatin. After vehicle or agonist stimulation, RNCMs were incubated for 1 hour at physiological conditions whereafter RNCMs were harvested, and RNA was isolated.

### Differentiation of human induced pluripotent stem cells into cardiomyocytes

The use of human derived hiPSCs in this research was approved by the McGill University Health Centre Research Ethics Board. In this paper, the hiPSC line derived from a healthy donor (AIW002-2) was provided from the Montreal Neurological Institute through the Open Biorepository, C-BIGR [17]. Human iPSCs, between passages 2 to 8 post-thawing, were differentiated into cardiomyocytes following the established GiWi protocol with slight modifications [18]. hiPSC-CMs were routinely screened for mycoplasma contamination and both iPSCs and differentiated iPSC-CMs used in this study were mycoplasma-free. For cardiomyocyte lineage commitment, a monolayer of 500,000 iPSCs, dissociated with Accutase, were seeded onto Matrigel (Corning, 354277) coated 24-well dishes in mTeSRPlus media supplemented with Rho kinase inhibitor, Y-27632-HCl (Selleckchem, S1049). The next day, the media was exchanged for fresh mTeSRPlus media. On day 0 of the protocol, iPS cells were stimulated with Wnt activator, CHIR99021 (Cayman Chemical, 13122) for 24 hours in RPMI 1640 media supplemented with B27 minus insulin (ThermoFisher, A1895602). At the same time the next day, the media was exchanged for fresh RPMI 1640 supplemented with B27 minus insulin. On day 3, Wnt was inhibited using the IWP2 compound (Selleckchem, S7085). The media was exchanged 48 hours later with RPMI 1640 supplemented with B27 minus insulin. On day 7 and day 10 of the protocol, the media was exchanged for RPMI 1640 supplemented with regular B27 (ThermoFisher, 17504001). Starting at day 12, metabolic selection was conducted where the cells were starved from glucose for 5-6 days. To promote cardiomyocyte survival and deplete undifferentiated cells, RPMI 1640 without glucose (Wisent, 350-060-CL), supplemented with B27 as well as with 4 mM lactate was used [19]. Wells that contained spontaneously beating cardiomyocytes were reseeded into 6-well dishes coated with fibronectin in PBS and maintained in RPMI 1640 media supplemented with B27 until day 28 where hiPSC-CMs were collected for RNA extraction. Prior to signaling experiments, hiPSC-CMs were seeded into optical bottom, black 96-well plates for cellular signalling experiments (Thermo Scientific, 165305).

### Cloning biosensors in AAV plasmids

The FRET-based ERK_1/2_ protein kinase biosensors EKAR-EV were generously provided by Dr. Michiyuki Matsuda and carried a nuclear localization (NLS) signal sequence [20]. This FRET-based biosensor was introduced into an AAV compatible backbone, pENN-AAV-CAG-tdTomato (Addgene catalog #105554) using a ‘cut-and-paste’ method using BamHI and BstBI restriction enzymes and ligase. The ExRai-AKAR2-NLS biosensor was obtained from Dr. Jin Zhang’s lab [21]. For cloning ExRai-AKAR2 biosensor into an AAV backbone, a multiple cloning site (MCS) was first generated in pAAV-CAG-hChR2-H134R-tdTomato (Addgene catalog #28017). The MCS was generated by annealing the following two primers 5’-gatccgctagcgtttaaacttaaggtaccgagctcactagtgaattctgcagatatccagcacagtggcggccgctcgagggcccttcg a-3’ and 3’-gcgatcgcaaatttgaattccatggctcgagtgatcacttaagacgtctataggtcgtgtcaccgccggcgagctcccgggaagcttcga-5’. To introduce the ExRai-AKAR2 biosensors within the pAAV-CAG-MCS backbone, biosensors were amplified by PCR and inserted using 5’-NheI and 3’-HindIII. Universal forward primer sequence was 5’-gctagctagcgccaccatgctgcgtcgcgccaccctg-3’. Reverse primer was used 5’-catagaagcttttatgcgtcttccacctttc-3’ for ExRai-AKAR2-NLS.

### Transduction of primary neonatal rat and iPSC-derived cardiomyocytes

Adeno-associated viruses (AAVs) used in this study were produced by the Neurophotonics Platform Viral Vector Core at Laval University, Québec, Canada. Upon arrival, AAVs were aliquoted in low-retention Eppendorf tubes to minimize freeze-thaw cycles. AAVs were stored for long-term in the −80°C and once thawed, were stored at 4°C for a maximum of 7-10 days. Cardiomyocyte cultures were transduced with biosensors packaged within AAV serotype 6 which is aligned with observations made by other groups demonstrating this serotypes effectiveness at infecting cardiomyocytes [22, 23]. During the transduction protocol, AAVs were kept on ice and diluted in maintenance media for each type of cardiomyocyte. Prior to transduction, cardiomyocyte cultures were washed three times to remove excess cell debris carried over during primary cell isolation or the hiPSC-CM re-seeding step. For effective infection, a multiplicity of infection 5000, indicative of 5000 virions per cell, 1.5 x 10^8^ virions per well, was used for all experiments. Both RNCM as well as hiPSC-CM, were transduced for 72 hours to allow for sufficient biosensor expression. This time frame was chosen as RNCMs have limited time in culture.

### High-content imaging of primary neonatal rat and hiPSC-derived cardiomyocytes

Prior to conducting the signalling experiments, cardiomyocyte cultures were inspected under a phase-contrast microscope to ensure RNCMs and hiPSC-CMs were healthy. Visual inspection of the cardiomyocytes permitted us to confirm that the hiPSC-CMs were spontaneously contracting and that both hiPSC-CMs and RNCMs appeared to have cardiomyocyte-*like* morphologies. In this study, the assay buffer was clear HBSS with calcium, magnesium, and sodium bicarbonate (Wisent, 311-513-CL). Sterile assay buffer was warmed to physiological temperature, 37°C, and cardiomyocytes were washed 2-3 times prior to imaging. Cells were left bathing in 90 μL of HBSS for imaging. Post-washing, cardiomyocyte cultures were re-incubated in a humidified atmosphere of 37°C with 5% CO_2_ for appropriately 1 hour before imaging to allow cells to re-equilibrate in the assay buffer. During this time, the temperature control settings (TCO) of Perkin Elmer’s Opera PHENIX high-content screening system were set allowing sufficient time for to warm up to 37°C with 3% CO_2_ for live-cell imaging. All drugs were prepared 10-fold more concentrated as 10 μL would be added in each well representative of a 10-fold dilution, 100 μL final volume. For nuclear ERK_1/2_ assays, images were acquired using a 20X air objective using a 425 nm laser for excitation of CFP. Emissions were detected with filters at 435-515 nm (CFP-donor) and 500-550 nm (YFP-acceptor). For the ExRai-AKAR2-NLS biosensor, images were acquired using a 20X air objective using a 375 nm and 480 nm laser for excitation and 500-550 nm emission filter. Within Perkin Elmer’s Harmony software, the experiment was set up in such a way where the imager would first acquire a ligand-independent measurement before the microwell plate would be ejected allowing time to perform drug stimulations. The plate itself was not displaced during drug treatment. Post-treatment measurements were automated and continued for another 70 minutes for a total of 8 readings, acquired at 10-min intervals.

### Image analysis and data processing in R

After live-cell imaging, images were imported into Perkin Elmer’s Columbus analysis software. Raw image files were also transferred onto external hard drives for long-term safe keeping. In Columbus, images were processed by first identifying the nuclei of individual cardiomyocytes followed by the calculation of morphology features. This output provided information regarding individual nuclei size and shape including roundness. To remove cell debris from the analysis, a size threshold was set as well as a roundness criterion. Intensity properties were calculated for both CFP and YFP fluorophores for EKAR-NLS and the same was done for the single fluorophore ExRai-AKAR2-NLS biosensor. Next, FRET- and excitation-ratios were computed by dividing acceptor/donor (YFP/CFP) for EKAR-NLS and 488 nm / 375 nm for the ExRai-AKAR2-NLS sensors, respectively. Columbus was then queried to output these data for each cardiomyocyte that fit the criteria listed and data was exported as text files for further analysis. Data were then processed in R based on a previously published in-house single-cell analytical approach with slight modifications, formerly applied to neuronal cultures [24]. Briefly, cardiomyocyte nuclei that appeared in all 8 timepoints were carried forward. As nuclei tend to drift slightly overtime, occasionally due to minor microplate positional changes occurring during ligand addition, we set a parameter where the fluorescence intensity of each nucleus could only deviate by ≤20% intensity to be considered as the same object/nuclei. To calculate the delta FRET or change in FRET in response to drug stimulations, we subtracted the ligand induced FRET from basal, ligand independent FRET. This expression was then converted into a percentage change in FRET (%ΔF/F) where the denominator (F) represented basal FRET. For consistency, (F) was computed by averaging the basal FRET across all nuclei in the same microwell. To detect whether cardiomyocyte sub-populations exhibited differential agonist-induced responses, when comparing RNCMs and hiPSC-CMs, we applied a clustering algorithm using the TSclust package [25] in R. Nuclei responses were clustered based on their magnitude of response over time, using the ‘pam’ function. For the single cell clustering, hiPSC-CMs and RNCMs nuclei were merged into a single dataset. Merging both sets of data allowed for output clusters to be reliable across RNCMs and hiPSC-CMs as means to obtain matched patterns and clusters within the two populations being compared. The algorithm could yield distinct clusters when applied to independent data sets. Following clustering, the merged hiPSC-CM and RNCM datasets were split and further plotted as heatmaps for visualization using pheatmap package. Stacked bar charts were also generated to better summarize the data using the ggplot2 package.

### RNAseq preparation and analysis

RNCM and hiPSC-CM cultures were seeded on fibronectin-coated 6-well dishes at a density of 1 million cells per well. After vehicle and drug stimulations, followed by 1 hour incubation, RNA was collected and isolated with the QiAshredder kit (Qiagen, 79656) followed by RNeasy™ Mini Kit (Qiagen, 74106) according to manufacturer’s instructions. RNA quality was later assessed by using a NanoDrop to ensure for RNA purity. Libraries were prepared using the NEBNext™ rRNA-depleted (HMR) stranded library kit and paired-end 100 bp sequencing to a depth of 25 million reads per sample. Sequencing was performed on the Illumina NovaSeq™ 6000 at the McGill University and Génome Québec Innovation Centre. RNAseq analysis was performed as we have previously described [15, 26]. Briefly, quality of reads was determined using FastQC, and trimmed with TrimGalore. Trimmed reads were then aligned to the Ensembl rat reference genome (Rattus_norvegicus.Rnor_6.0.100) or the reference human genome (Homo_sapiens.GRCh38.104) with STAR. Transcripts were then assembled with StringTie. Normalized transcript abundance, measured as TPM, transcript per million, was plotted using pheatmap package in R. Curated gene sets were downloaded from https://www.gsea-msigdb.org/gsea/index.jsp.

### Statistical analysis

All statistical analyses were performed using GraphPad Prism software where summary data processed and computed in R was imported. For cellular signalling datasets, Figures 2, 3 and 4 and supplementary Figure 4, non-parametric Welch’s t-test were conducted as test statistic to determine whether the RNCMs and hiPSC-CM populations and clusters had equal means. In supplemental Figure 3, we performed Welch’s ANOVA followed by Dunnett’s multiple comparisons test, to test whether the response means were equal across different fields measured. Welch’s ANOVA was chosen as Bartlett’s test statistics indicated that there was a significant difference between the variances.

**Figure 1.**
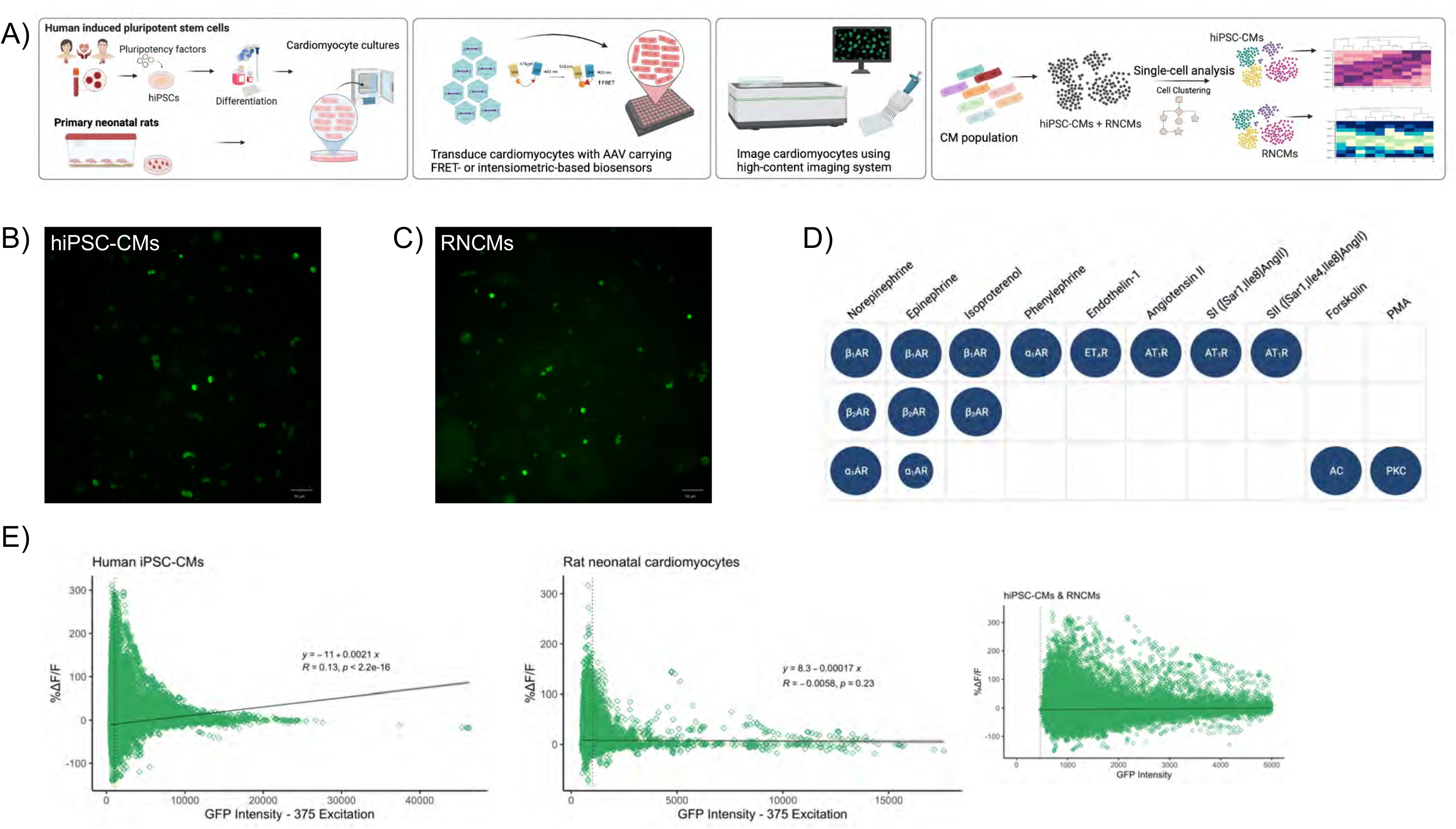
Measuring protein kinase activity in RNCMs and hiPSC-CMs. **(A) Depiction of the experimental pipeline used to compare signalling signatures in two cardiomyocyte model systems.** The experimental pipeline begins with the differentiation of hiPSCs into cardiomyocytes or the isolation of rat neonatal cardiomyocytes. Cardiomyocyte cultures are then transduced with adeno-associated virus serotype 6 (AAV6) to introduce the biosensor of interest. Next, RNCM and hiPSC-CM cultures are imaged using high content microscopy and further analyzed using a single cell analytical approach. Representative fluorescent microscopy images illustrating the expression of ExRai-AKAR2-NLS biosensor in **(B)** hiPSC-CMs and **(C)** RNCMs. **(D) Diagram depicting the drugs used in this study.** After a baseline reading was taken, RNCMs and hiPSC-CMs were stimulated with saturating doses of a panel of ligands that targeted cardiac relevant GPCRs. Norepinephrine, epinephrine, isoproterenol, phenylephrine, SI and SII were used at 10 μM. Ang II and PMA were used at 1 μM. Forskolin was prepared as 5 μM and ET-1 at 100 nM. **(E) Basal, ligand independent single fluorophore intensities of hiPSC-CM and RNCM ExRai-AAKR2-NLS datasets.** Biosensors were expressed at higher levels in hiPSC-CMs compared to RNCMs despite being transduced for the same length of time.

**Figure 2.**
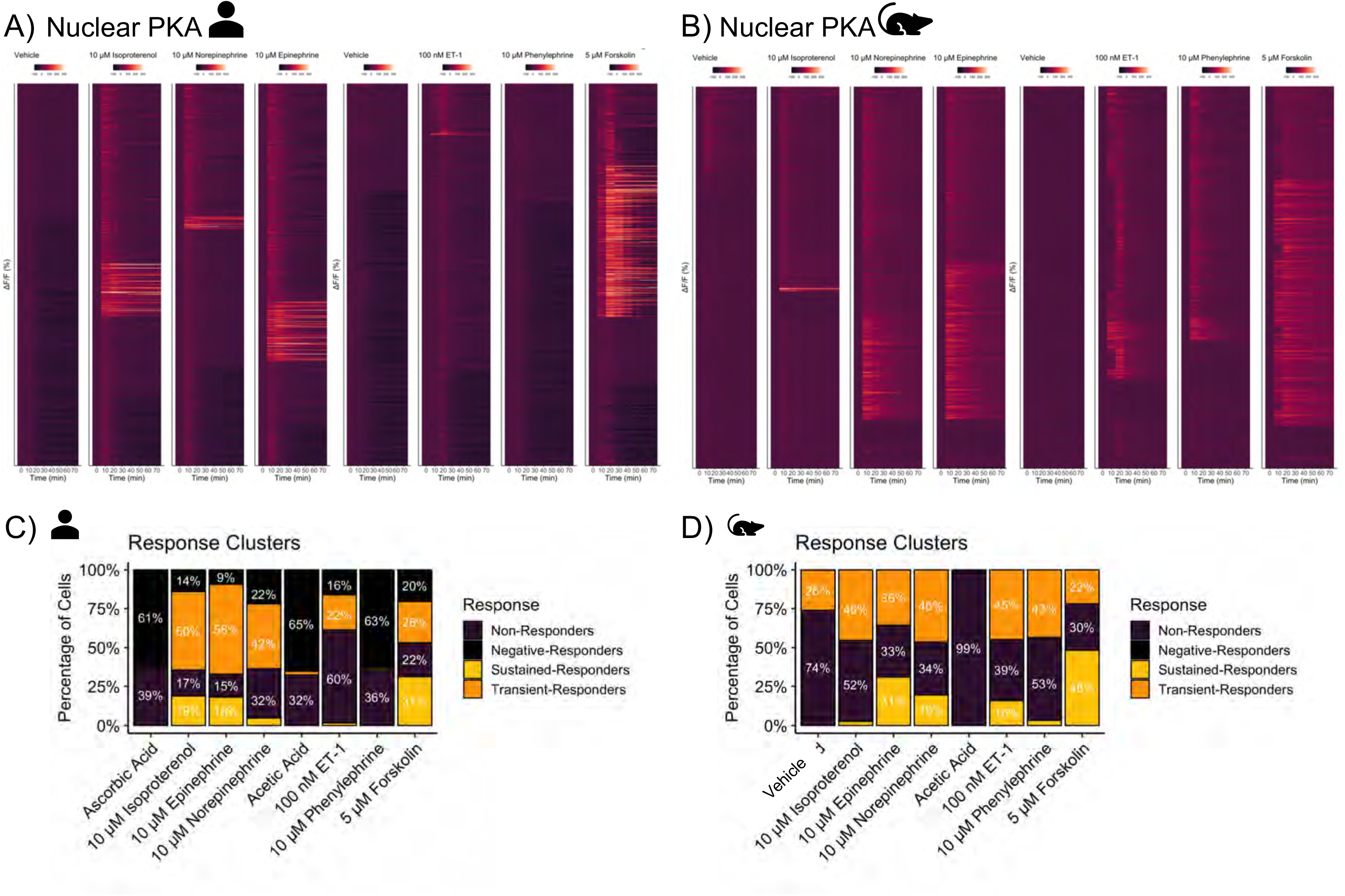
Single nuclei PKA activation patterns as observed in hiPSC-CM and RNCM cultures in response to adrenergic agonists and vasoactive ET-1 peptide. Heatmaps displaying nuclear PKA activity as measured in **(A)** hiPSC-CMs and **(B)** RNCMs. Single nuclear PKA data summarized as %ΔF/F (y-axis) as a function of time (x-axis). The data was partitioned into four clusters representing distinct nuclear behaviors, either exhibited sustained or transient responses to agonists while other nuclei failed to respond or experienced a decrease in activity compared to baseline. Representation of the four response clusters was plotted as a bar chart with percentages of nuclei belonging to each response cluster as observed in **(C)** hiPSC-CMs and **(D)** RNCMs. Experiments were performed using RNCMs isolated from 4 different neonatal rat pup litters and 3 independent cardiac differentiations.

**Figure 3.**
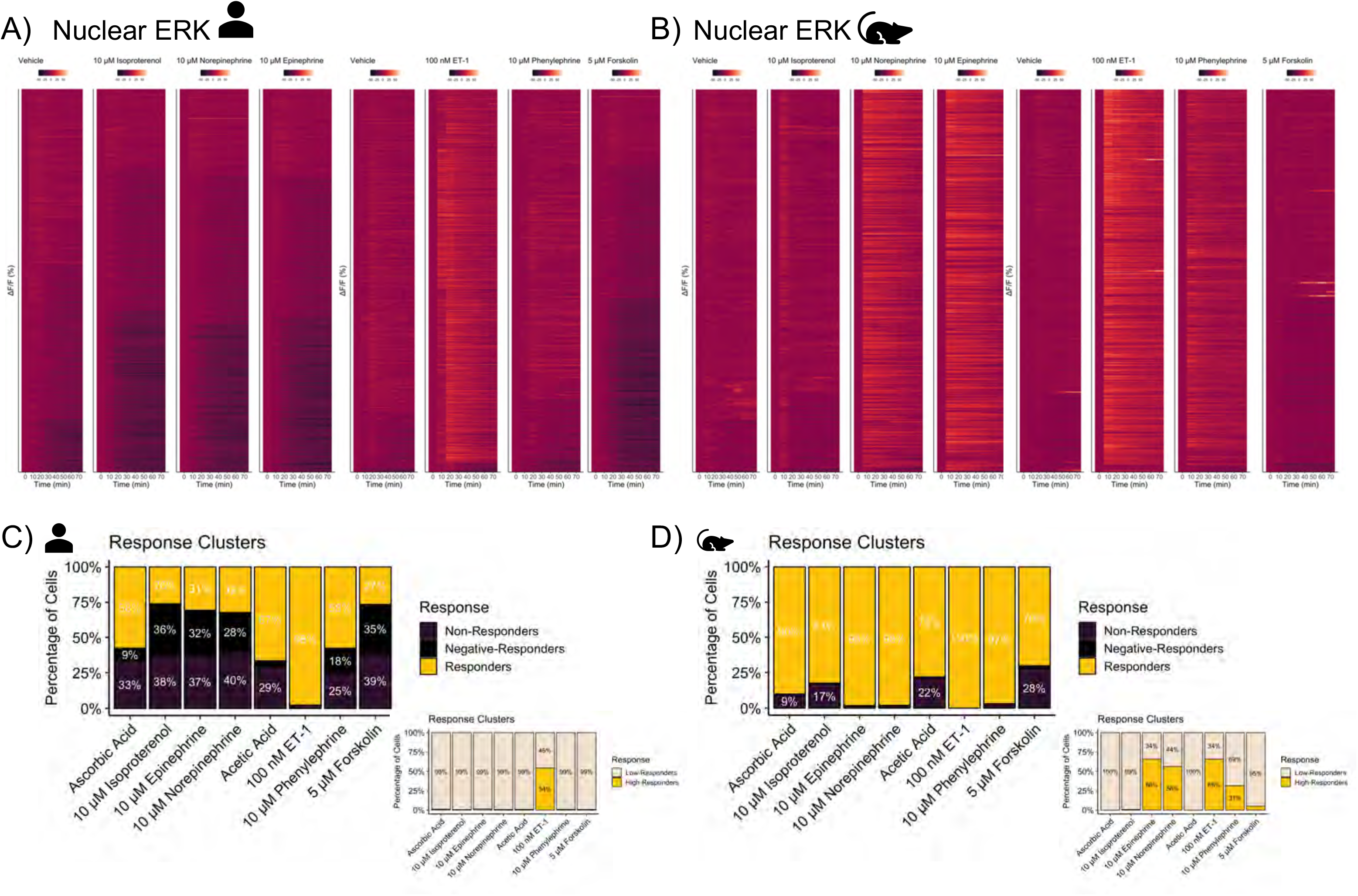
Single nuclei ERK_1/2_ activation patterns as observed in hiPSC-CM and RNCM cultures in response to adrenergic agonists and vasoactive ET-1 peptide. Heatmaps displaying nuclear ERK_1/2_ activity as measured in **(A)** hiPSC-CMs and **(B)** RNCMs. Single nuclear ERK_1/2_ data summarized as %ΔF/F (y-axis) as a function of time (x-axis). The data was partitioned into three clusters representing distinct nuclear behaviors. CM nuclei either responded to the drug stimulations, resulting in ERK_1/2_ activation, while others did not respond, and the last cluster of nuclei represented those that exhibited a decrease in ERK_1/2_ activity compared to baseline. The three response clusters were plotted as stacked bar charts with percentages of nuclei belonging to each cluster as observed in **(C)** hiPSC-CMs and **(D)** RNCMs. *Insets* reflect sub-clustering applied to the ‘responding’ nuclei cluster. It is apparent that within the ‘responding’ sub-population, two other clusters can be identified based on their responding magnitudes-low or high responders. Experiments were performed using RNCMs isolated from 3 different neonatal rat pup litters and 3 independent cardiac differentiations.

**Figure 4.**
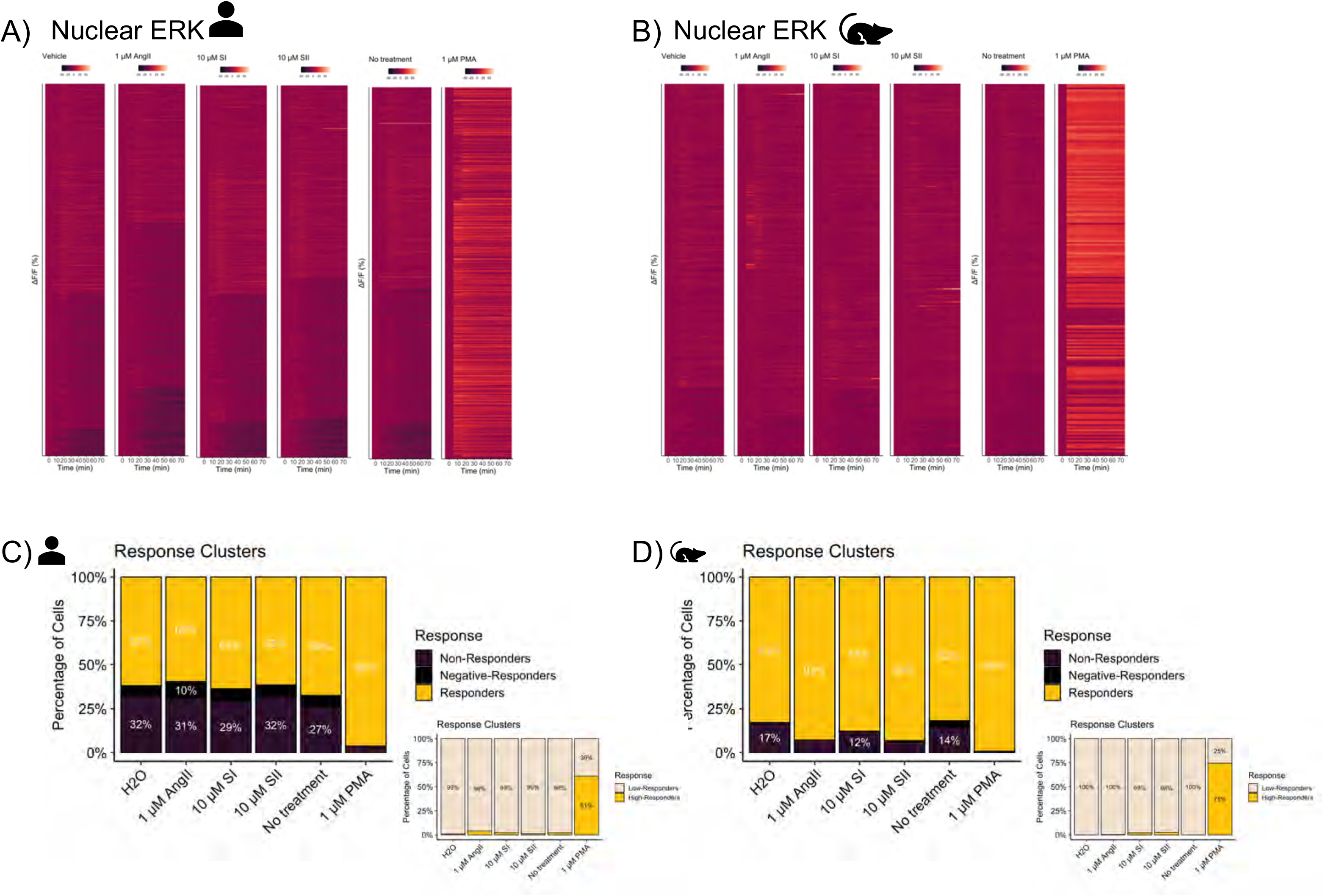
Single nuclei ERK_1/2_ activation patterns as observed in hiPSC-CM and RNCM cultures in response to Ang II and β-arrestin biased peptides. Heatmaps displaying nuclear ERK_1/2_ activity as measured in **(A)** hiPSC-CMs and **(B)** RNCMs. Single nuclear ERK_1/2_ data summarized as %ΔF/F (y-axis) as a function of time (x-axis). The data was partitioned into three clusters representing distinct nuclear behaviors. CM nuclei either responded to the drug stimulations, resulting in ERK_1/2_ activation, while others did not respond, and the last cluster of nuclei represented those that exhibited a decrease in ERK_1/2_ activity compared to baseline. The three response clusters were plotted as stacked bar charts with percentages of nuclei belonging to each cluster as observed in **(C)** hiPSC-CMs and **(D)** RNCMs. *Insets* reflect the sub-clustering applied on the ‘responding’ nuclei cluster. It is apparent that within the ‘responding’ sub-population, two other clusters can be identified based on their responding magnitudes-low or high responders. Experiments were performed using RNCMs isolated from 3 different neonatal rat pup litters and 3 independent cardiac differentiations.

## Results and Discussion

### Distinct protein kinase activation patterns noted in hiPSC-CM and RNCM cultures

#### Nuclear PKA signalling

To compare the cellular contexts of hiPSC-CMs against rat neonatal cardiomyocytes, we began probing for differences in cellular signalling. Here, we set out to test the null hypothesis that RNCMs and hiPSC-CMs are equivalent cellular models. With this in mind, we measured kinase activation in RNCMs and hiPSC-CMs, at the single cell level, every 10 minutes for 70-minutes, as in the experimental pipeline depicted in **Fig 1A**. To probe agonist-induced PKA activation profiles, cardiomyocyte cultures were transduced with AAV2/6-ExRai-AKAR2-NLS (**Fig 1B, C**). A few days later, cardiomyocyte cultures were imaged using an automated high content imaging system. Baseline measurements were recorded prior to stimulating cells with a panel of saturating doses of GPCR agonists targeting either endothelin, adrenergic or angiotensin systems. (**Fig. 1D**). As seen in **Fig. 1E**, the expression level of the nuclear-localized PKA biosensor, as represented on the x-axis, was considerably higher in hiPSC-CMs compared to RNCMs, despite being transduced for the same length of time. When testing the relationship between biosensor expression (RFU) and biosensor output (%ΔF/F), we did not observe a significant linear correlation in the RNCM cultures as shown with small R correlation coefficient and nonsignificant p-value (p =0.23). In the hiPSC-CMs, the R coefficient was small, yet the p-value was significant, an observation that is suggesting greater variability around the regression line and a larger prediction interval when correlating both variables. Thus, to have a more uniform population of RNCMs and hiPSC-CMs, removing plausible confounds caused by biosensor expression, we set a threshold and analysed CMs on the lower end of the spectrum. Cardiomyocytes that expressed the PKA sensor between 0 and 5000 RFU were carried forward, and no cells exhibited zero intensity as the lowest unit measured was 462 RFU (**Fig. 1E**, inset).

Next, we averaged the nuclear PKA responses of all CMs, across our biological replicates, and plotted the responses as a function of time (**Supp. Fig. 1**). RNCMs and hiPSC-CMs displayed distinct averaged PKA signatures when stimulated by various agonists, when evaluating both kinetics and magnitude of the responses. In both CM types, forskolin, a direct activator of adenylyl cyclase, was the most potent activator of the PKA pathway followed by epinephrine. Markedly, epinephrine and norepinephrine led to larger nuclear PKA responses in RNCMs. Phenylephrine and ET-1 also resulted in PKA activation; an observation examined later. To move beyond averaged responses, and as means to fully appreciate subtle differences or granularity between RNCMs and hiPSC-CMs, we applied a single cell analytical approach to our biosensor datasets [24]. To identify cardiomyocyte populations or clusters that responded differently within the overall population, we merged RNCM and hiPSC-CM datasets and ran them through a clustering algorithm [25]. The clustering technique revealed that the overall population of cardiomyocytes could be sub-divided into 4 main clusters (**Suppl. Fig. 2**). We arbitrarily named them, ‘sustained-responders’, ‘transient-responders’, ‘non-responders’ and ‘negative-responders’. To test whether the expression of the biosensor influenced the clustering method, cardiomyocytes at different fluorescent intensities thresholds were independently ran through the algorithm (**Supp. Fig. 2A**). The inclusion of cells at varying fluorophore intensities did not appreciably impact how the cells were clustered. We also noticed that while requesting the clustering algorithm to devise four clusters instead of three (**Supp. Fig. 2B**), the data was partitioned into *more* informative patterns. By requesting four clusters, we saw that the population of ‘responders’ could be further sub-divided based on kinetics, with some nuclei exhibiting ‘transient’ while other displayed ‘sustained’ effects. Four clusters were carried forward throughout further analyses. As another internal control, we investigated whether all cardiomyocytes seeded in the microwell were exposed to a uniform concentration of drug. When stimulating the cells, we were vigilant to break the surface tension barrier with the microtip to ensure effective drug diffusion, as we were working with small, 10 μL volumes. The 21 fields imaged were plotted against the percentage change in response compared to baseline, %ΔF/F, post-drug stimulation (**Suppl. Fig. 3**). No significant differences were observed in response to stimulation with norepinephrine while responses to forskolin and isoproterenol showed minor positional effects by field. Still, these observations may just as likely be correlated with the proportion of nuclei that fall within a response cluster, irrespective of drug diffusion. Thus, the fraction of nuclei exhibiting distinct behaviors, categorized as non-responders or responders may be skewing the average response per field.

To visualize the overall spread of single nuclei PKA responses, we built heatmaps with single nuclei responses plotted on the y-axis as a function of time (x-axis). In hiPSC-CMs, adrenergic targeting ligands, isoproterenol, norepinephrine, and epinephrine, resulted in modest activation of nuclear PKA (**Fig. 2A**). More specifically, the distribution revealed that 18.72 ± 5.73 % of hiPSC-CMs exhibited a sustained response to isoproterenol compared to 2.66 ± 1.5 % of RNCMs (**Fig. 2B**). Further, in response to epinephrine, 57.83 ± 2.3 % of hiPSC-CMs led to a transient response compared to 35.51 ± 13.5 % of RNCMs. Forskolin steered a similar response profile in both species with slightly more sustained responders in the overall RNCM population. Most remarkably, phenylephrine resulted in nuclear PKA activity in RNCMs but not in hiPSC-CMs (**Fig. 2B**). A greater density of α_1_AR in RNCMs could explain the prominent PKA response upon stimulation with phenylephrine. Similarly, ET-1 drove a transient PKA activation profile in RNCMs and hiPSC-CM, albeit to a smaller degree in iPSC-CMs. As the ET_A_ receptor is canonically coupled to Gα_q_, this observation was unpredicted. However, this finding has been previously observed in both HeLa cells overexpressing the ET_A_ as well as in rat aortic smooth muscle cells [27]. In these models, PKA activation was reported to be independent of cAMP production. Further, in rat aortic vascular smooth muscle cells, ET-1 coupling to Gα_i_ was shown to result in transient PKA activation, with a return to baseline after 20 minutes [28]. An observation analogous to what we observed in RNCM cultures with approximately 45 ±15 % of cells exhibiting this transient behaviour (**Fig. 2B**). G proteins may be expressed at varying stoichiometries in both cell types, examined in a later section. However, this observation contrasts with one of our previous reports which demonstrated phenylephrine’s ability to induce nuclear PKA responses in RNCMs but not ET-1 [15]. However, as ExRai-AKAR2-NLS is more sensitive than the AKAR2-NLS sensors used in our previous study, this discrepancy may simply reflect differential affinity and sensitivities of the biosensors themselves. Nonetheless, this result is worth follow-up studies as it may be implicated in shaping transcriptional programs involved in the development of cardiac hypertrophy [15]. With a greater proportion of hiPSC-CMs exhibiting a decrease in PKA activity compared to baseline, this behaviour may suggest that adrenergic receptors exhibit a greater degree of constitutive activity in this context. Lastly and as expected, compounds that targeted the AT_1_R did not result in PKA activation (**Suppl. Fig. 4**). Interestingly, Ang II and related compounds led to a measurable negative response in hiPSC-CMs. PMA, a direct activator of PKC, also failed to trigger PKA activation, except for a small population of hiPSC-CM. This may reflect signalling cross-talk [29]. Altogether, there are trends that indicate distinct signalling landscapes in both species.

#### 3.1.2 Nuclear ERK_1/2_ signalling

We were also interested in probing for activation of extracellular signal-regulated kinase, ERK_1/2_. Correspondingly, RNCMs and hiPSC-CMs were transduced with AAV2/6-EKAR-NLS to measure nuclear ERK_1/2_ activity (**Suppl. Fig. 5A, B**). As observed in our PKA experiments, hiPSC-CMs expressed the EKAR-NLS biosensor at higher donor, CFP intensities compared to RNCMs (**Suppl. Fig. 5C**). With our available computer memory (RAM), it was not possible to include all CMs as too many objects within R clustering exhausted the memory limit. Therefore, for consistency in our data analysis, we filtered our CM populations and carried forward CMs that expressed the biosensor between 2000 and 5000 RFU. The response clusters did not appear to change greatly with higher CFP intensities except at higher CFP intensities as less RNCMs fit that criterion (**data not shown**). However, before dissecting single cell response profiles, we plotted averaged ERK_1/2_ responses over time. The summarized data demonstrated that PMA and ET-1 were the most robust responders in RNCMs and hiPSC-CMs, displaying the greatest response magnitudes, while epinephrine, norepinephrine and phenylephrine exhibited larger ERK_1/2_ activation in RNCMs compared to hiPSC-CMs (**Suppl. Fig. 5D**).

Next, we plotted the single cell ERK_1/2_ responses using heatmaps (**Fig. 3, 4**). In hiPSC-CM profiles, adrenergic ligands displayed considerable variability, ~30% of nuclei that exhibited a decrease in ERK_1/2_ activation compared to baseline. Adrenergic receptors may be exhibiting a greater degree of constitutive activity in this context, or this observation may expose a subpopulation of cells signalling through a β_2_AR-Gα_i_ pathway. In hiPSC-CMs, ET-1 treatment led to the most robust and sustained response over time in 97.8 ±0.5% of the nuclei population. However, even vehicle showed ERK_1/2_ activity albeit to a considerably smaller magnitude. To further dissect this response range, low and high responses, we applied the clustering algorithm on a select population, those previously categorized as ‘responders’. This yielded two more subclusters, arbitrarily named ‘low-responders’ and ‘high-responders’ (**Suppl. Fig. 6C**). This extra step allowed us to summarize the data in a way that better reflected the spread as observed with the heatmaps (**Fig. 3 C, D inset**).

In hiPSC-CMs, phenylephrine effects were less pronounced but not significantly stronger than vehicle (**Fig. 3A**). In strike contrast, adrenergic ligands resulted in significant nuclear ERK_1/2_ activity, specifically in response to epinephrine (p=0.033) and norepinephrine (p=0.038) in RNCMs compared to hiPSC-CMs (**Fig. 3B, D inset**). In RNCMs, transient activation patterns were observed in response to isoproterenol (p= 0.047), an observation that was not mirrored in hiPSC-CMs. Phenylephrine drove ERK_1/2_ activation to a greater extent in RNCMs compared to hiPSC-CMs, once again suggesting that hiPSC-CM and RNCMs may express α_1_ARs at different levels. In adult mouse cardiomyocyte cultures, phenylephrine has been shown to result in ERK activation in 60% of CMs, as α_1_AR receptors have been demonstrated to be expressed in 60% of ventricular cardiomyocytes [30, 31]. This observation is similar to our ERK_1/2_ assays conducted in hiPSC-CMs, at 58% (**Fig. 3C**). This finding is aligned with reports suggesting that mouse CM cultures are better mimics of the human CM, when studying α_1_AR-mediated myocardial effects, as α_1_AR expression levels are more comparable. In contrast, rats express 5-10-fold more α_1_AR, a finding that is aligned with the strong phenylephrine responses observed in our RNCM experiments [32, 33] (**Fig. 3 B, D**).

Unsurprisingly, forskolin failed to produce significant ERK_1/2_ signals in either CM models while PMA drove activation in 99% of CMs, irrespective of the species. Similarly, the Ang II peptide and β-arrestin biased drugs, SI and SII did not drive significant ERK_1/2_ activity in comparison to the vehicle (**Fig. 4C, D inset**). It is possible that by the time our cells were imaged, at the 10-minute time point, the early ERK_1/2_ activation phase has passed as we have data western blot data that supports ERK_1/2_ activity as early as 5 minutes in RNCMs (**Suppl. Fig. 7**). However, in the abovementioned single cell FRET study, the authors also reported the lack of an Ang II-mediated ERK response.

### Bulk RNA-seq reveals transcriptome-level differences in three cells types: hiPSC-CMs, RNCMs and HEK 293 cells

#### GPCR-mediated signal transduction-related gene sets

As distinct signalling signatures were observed at the level of protein kinase activation, we next sought to test whether these differences could be explained via mRNA expression analyses. Thus, we set out to determine if RNCM and hiPSC-CMs were wired differently, at the endogenous level. To characterize the transcriptional landscapes in hiPSC-CM and RNCM function, we surveyed genes linked with GPCR signalling at baseline, independent of ligand. Since HEK 293 cells have been a valuable model system for dissecting GPCR functions *in vitro*, we included these cells in our analysis.

Basal abundance of select cardiac-relevant Class A GPCRs, reported as transcript per kilobase million, TPM, revealed striking differences between the three *in vitro* cell models assessed (**Fig. 5**). As suggested earlier, indeed, the α_1_-adrenergic receptor was more highly expressed in RNCM compared to hiPSC-CMs and HEK 293 cells (**Fig. 5A, Supp. Fig. 8A**). The β_1_-adrenergic receptor was also more abundant in RNCMs. The endothelin A (ET_A_) receptor was expressed at higher levels in hiPSC-CMs compared to RNCMs. In contrast, angiotensin II type I (AT_1_) receptors were expressed at low amounts in all three cell types. In addition, certain features of cardiac remodelling associated with the angiotensin system have been reported to occur through fibroblast activation and paracrine mechanisms [34]. Regarding the endogenous signalling machinery, including heterotrimeric G proteins, Gα_s_ was shown to be well expressed in all three cell types (**Fig. 5B, Supp. Fig. 8C**). Gα_q_ was demonstrated to be expressed at comparable levels in all three cell types, though at slightly higher levels in RNCMs. Gα_i2_ expression was reported as 3-fold higher in RNCMs. Conceivably, this may be associated with the PKA response to ET-1, as observed in **Fig. 2B**. Gβ isoforms were all observed, albeit to varying degrees, with Gβ1 being most abundant followed by Gβ2 (**Fig. 5C, Supp. Fig. 8D**). Equally, all Gγ isoforms were detected with Gγ5, 10 and 12 being most abundant (**Fig. 5D, Supp. Fig. 8E**). Lastly, distinct expression patterns were observed for select GPCR effector molecules (**Fig. 5E, Supp. Fig. 8F**). For example, adenylyl cyclase isoforms 5 and 6 were more abundant in cardiomyocytes compared to HEK 293 cells, an observation consistent with previous reports [35]. In contrast, β-arrestin2 was expressed at greater levels in HEK 293 cells, perhaps affecting GPCR desensitization or post G protein signalling. PKA enzyme subunits were well represented as well as members of the MAPK cascade, albeit at different level (**Fig. 5E, Suppl. 8F**). All in all, mRNA levels seem to align well with the cellular signalling signatures reported above. Further, differential DESeq2 analysis demonstrated that genes that were either significantly up- or down regulated in RNCMs and hiPSC-CMs after 60-min agonist stimulation differed (**Suppl. Fig. 9**). A 60-min time point was selected to parallel the time course of our signalling experiments. This finding indicates that their transcriptional landscapes are also different, driving distinct cellular effects. Taken together, evidence presented implies that cell context is a critical determinant dictating receptor outcomes. For physiologically relevant conclusions to be drawn, it is important to consider the cell context and its influence especially when translating research findings for human predictability [36].

**Figure 5.**
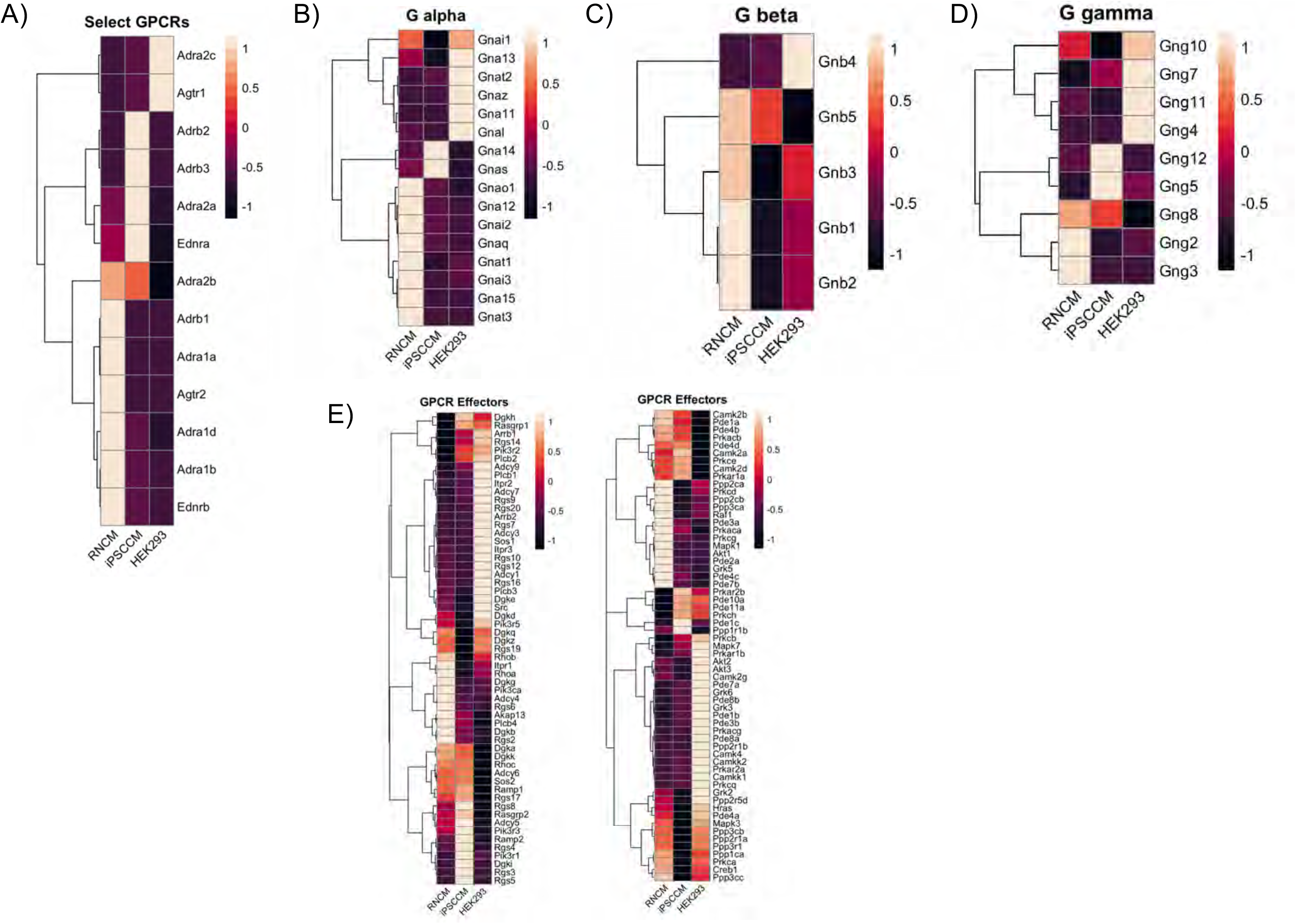
Ligand-independent RNA-seq based comparison of the endogenous signaling machinery expressed in RNCMs, hiPSC-CMs and HEK 293 cells. Exploratory analysis of gene sets associated with GPCR signal transduction investigating **(A)** select cardiac relevant class A GPCRs, heterotrimeric G proteins **(B)** Gα, **(C)** Gβ and **(D)** Gγ as well as **(E)** effector expression profiles. Colored heatmaps display comparative expression relative to all three cell models being assessed normalized as TPM. For comparison’s sake, data from HEK 293 cells was included (published originally as Lukasheva et al., 2020 Sci Rep 10, 8779. https://doi.org/10.1038/s41598-020-65636-3.

#### Myocyte-, maturity- and metabolism-related gene sets in hiPSC and RNCMs

We next examined gene sets associated with cardiomyocyte fate, maturity and ‘adult-like’ identity. When observing normalized transcript abundance of a curated gene list for cardiac progenitor differentiation, RNCMs as well as iPSC-CMs expressed GATA4, Nkx2-5 and Mef2c, key transcription factors that control cardiomyocyte fate (**Fig. 6A**). Sarcomeric proteins were also abundant including TNNT2, TNNi3, Actc1 and MYH6. In our cultures, the fetal troponin isoform, TNNI1 was more abundant in hiPSC-CMs compared to the adult equivalent, TNNI3 (**Fig. 6B**). RNCMs express approximately 5-fold more TTNI3 and 2.5-fold TNNI1 compared to day 28 hiPSC-CMs. This data is consistent with the current literature equating hiPSC-CMs as neonatal as opposed to adult cardiomyocytes. Both hiPSC-CMs and RNCM express MYH6, present in the developing ventricle and neither expressed the adult isoform MYH7 (**Fig. 6C**). This echoes to the neonatal nature of both of these cardiomyocyte cell models. Genes involved in glycolytic metabolism, such as ALDOA, BPGM, ENO1 and LDHA were more abundant in RNCM compared to hiPSC-CMs (**Suppl. Fig. 8A**) [37]. These genes have been shown to be downregulated in ‘matured’ hiPSC-CMs that undergo a metabolic switch from glycolysis to fatty acid oxidation. Overall, transcript reads, measured as TPM, were higher in glycolytic metabolism pathways compared to gene sets of fatty acid related metabolism. This result is suggestive that both RNCM and iPSC-CMs use glycolysis as an energy source.

**Figure 6.**
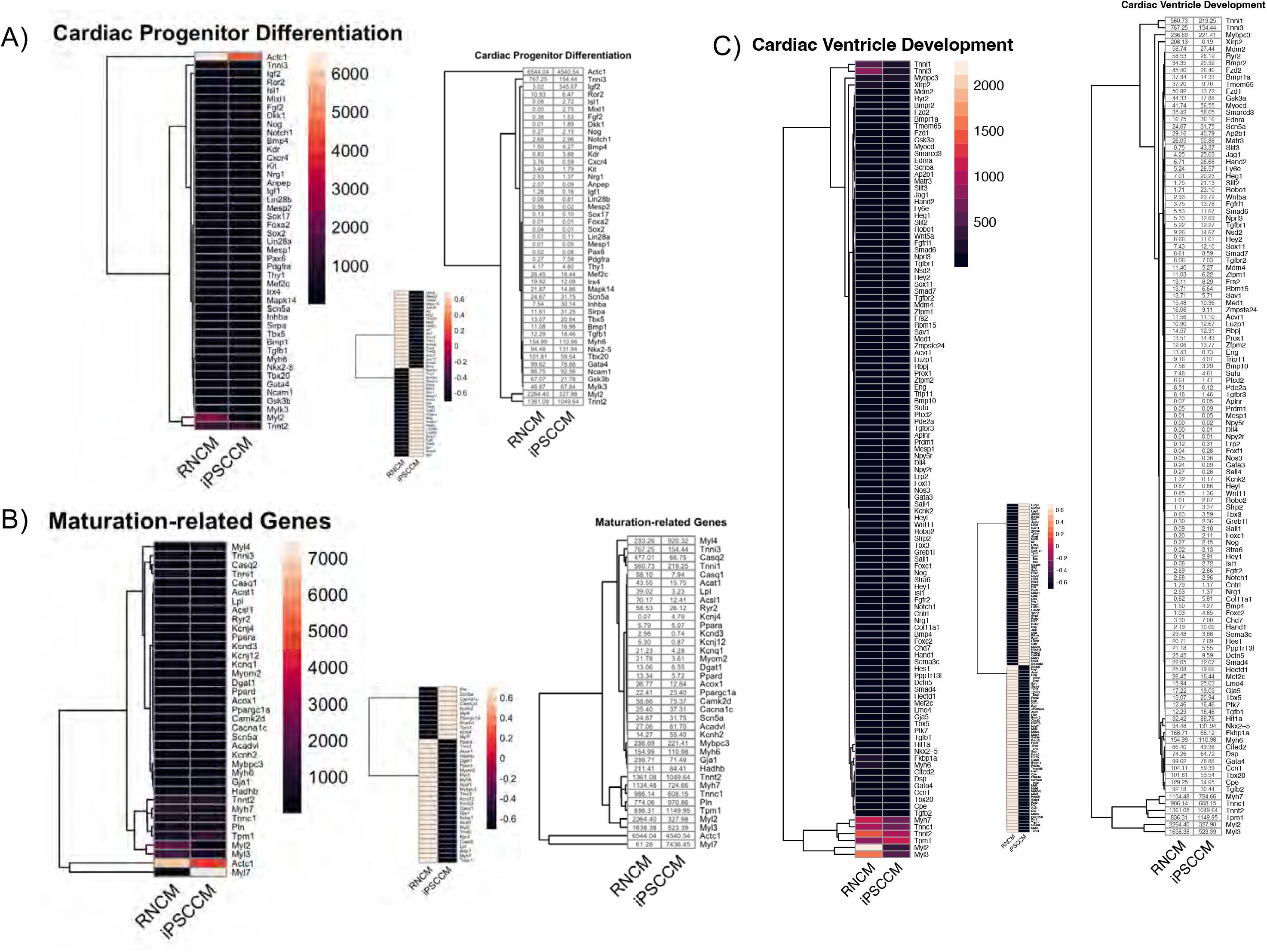
Ligand-independent comparative gene expression analysis in hiPSC-CMs and RNCMs. RNA-seq inferred transcript abundance measured as TPM of curated gene sets associated with **(A)** cardiac progenitor differentiation, **(B)** maturation-related genes and **(C)** cardiac ventricular development. Large colored heat maps represent gene expression based on TPM while smaller colored heatmaps display comparative expression (hiPSC-CM vs RNCMs). Colorless heatmap depict raw transcript abundance.

In the context of cardiomyocyte functionality and calcium regulation, both RNCM and iPSC-CMs express key genes: RyR2, Cacna1c, Camk2d, Atp2a2 (SERCA2), Casq2, Slc8a1 (NCX1) and Tmp1 (**Fig. 7A**). Transcript levels for Bin1, a gene that participates in T-tubule formation was low in both RNCM and iPSC-CMs, an observation consistent with the literature (**Fig. 7A, B**). In connection with genes involved in propagating the cardiac action potential, the voltage gated potassium channel, KCNH2 (hERG), was more abundant in hiPSC-CMs than RNCMs (**Fig. 7C**). The HCN4 subunit of the funny current, I_f_, was shown to be expressed at 2-fold higher abundance in hiPSC-CMs compared to RNCMs. This observation is in line with the automaticity seen in iPSC-CMs cultures and not in RNCMs [38]. The low (negligible: 0.35) transcript levels of KCNJ2 encoding for the inward rectifier potassium current *I_K1_* has also been associated with the spontaneous contractile properties of hiPSC-CMs [39]. All in all, our hiPSC-CMs display similar gene expression patterns as hiPSC-CMs cultures previously published and are similar to RNCMs in terms of maturity.

**Figure 7.**
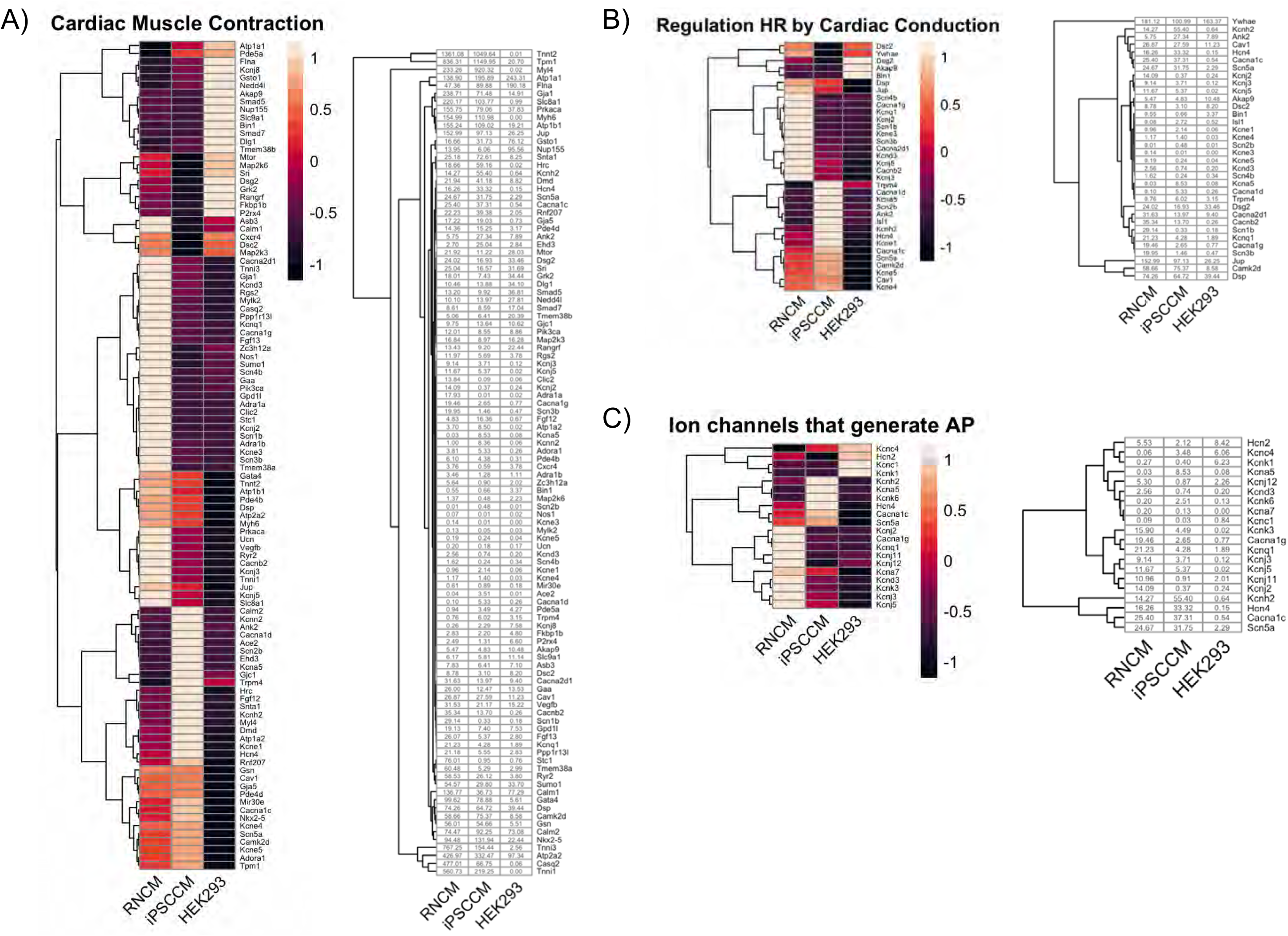
Gene-level examination of get sets associated with cardiomyocyte behavior and function. RNA-seq deduced transcript abundance of genes related with **(A)** cardiac muscle contraction **(B)** regulation of heart rate by cardiac conduction and **(C)** ion channels that generate cardiac action potential. Large colored heat maps represent gene expression based on TPM while smaller colored heatmaps display comparative expression (hiPSC-CM vs RNCMs). Colorless heatmap depict raw transcript abundance. For comparison’s sake, data from HEK 293 cells was included (published originally as Lukasheva et al., 2020 Sci Rep 10, 8779. https://doi.org/10.1038/s41598-020-65636-3.

#### Concluding remarks

Translational research programs take advantage of numerous immortalized, primary cell and now iPSC-derived cellular models to mimic the human condition. Several reports have attested to the usefulness of hiPSCs and their differentiated derivatives as human model systems especially when attempting to recapitulate disease *in a dish*. Still, we deemed it necessary to perform a comparative study with primary rat cardiomyocytes as hiPSC-CMs are associated with certain limitations, i.e., their inherent immaturity. As such, we combined single cell signalling with bulk transcriptomics to compare the traditional rat neonatal cardiomyocyte with human iPSC-CMs. Overall, our data serves to suggest that RNCMs and hiPSC-CMs exhibit unique signalling signatures that are, in part, rationalized by mRNA levels and cross-species differences. As these differences were captured in non-matured hiPSC-CMs, these observations point towards the notion that even if our hiPSC-CMs are not *adult-like*, they are still superior cellular *human* proxies compared to traditional rat primary cardiomyocytes.

Selecting the correct model system becomes even more relevant in the context of cardiovascular disease modelling as receptor and effector expression levels have been reported to be altered. In the human failing heart, for example, the ratios of β-adrenergic receptors, G protein, GRKs, are different compared to non-failing hearts [40]. These changes undoubtedly affect cellular signalling profiles, rewriting gene transcription. Therefore, as working with hiPSC-CM enables patient access, these model systems allow us to gain a glimpse at how cellular context affects signalling in disease. As hiPSC-CMs endogenously express the main therapeutic targets in heart failure, there is no need for the overexpression of recombinant proteins thereby generating undesired artificial systems, as commonly done in HEK 293 cells. Even if RNCM express the relevant players and effectors, their stoichiometries are different. Since their limited time in culture poses an extra experimental constraint, hiPSC-CMs represent an excellent proxy for modelling the human healthy or diseased heart in an *in vitro* setting. Lastly, hiPSC-CMs also express the hERG channel making them ideal models for cardiotoxicity studies as all drugs need to demonstrate the lack of pro-arrhythmic risk prior to approval.

This study does have potential limitations. It is important to consider certain confounding variables when analysing our single cell datasets. The rat neonatal cultures were essentially in an *in vivo* context 5 days prior to the assay and data collection while iPSC-CMs were in an *in vitro* context for 40-46 before being assayed. We attempted to control for this by obtaining rat iPSCs however the cells did not survive post-thawing. Secondly, even if the RNCM maintenance media was supplemented with a mitotic inhibitor, it is noteworthy to be aware that a small population of cardiac fibroblasts renders the overall population somewhat *impure*. However, our culturing method has been shown to yield a population composed of ≥90% cardiomyocytes. Besides, the differentiation protocol used in this study does yield a mixed population of ventricular, atrial and pacemaker cardiomyocytes. Thus, some of the clusters observed may reflect these different cell types as they do exhibit distinct behaviours. Thirdly, iPSC-CMs were not matured in this study, and it would be interesting to follow up and assess how maturated iPSC-CMs would compare to those used here as well as to RNCMs or adult primary cultures. Nevertheless, the main message of our study remains, that the cellular vehicle needs to express the correct stoichiometry of receptors and effectors for meaningful conclusions to be drawn from the data particularly for translation efforts. All in all, our study demonstrates that human stem cell derived models of the cardiomyocyte do provide significant advantages, even if immature, and should be taken advantage of if their use is appropriate and aligned with the research objective.

## Figure Legends

**Supplemental Figure 1.**
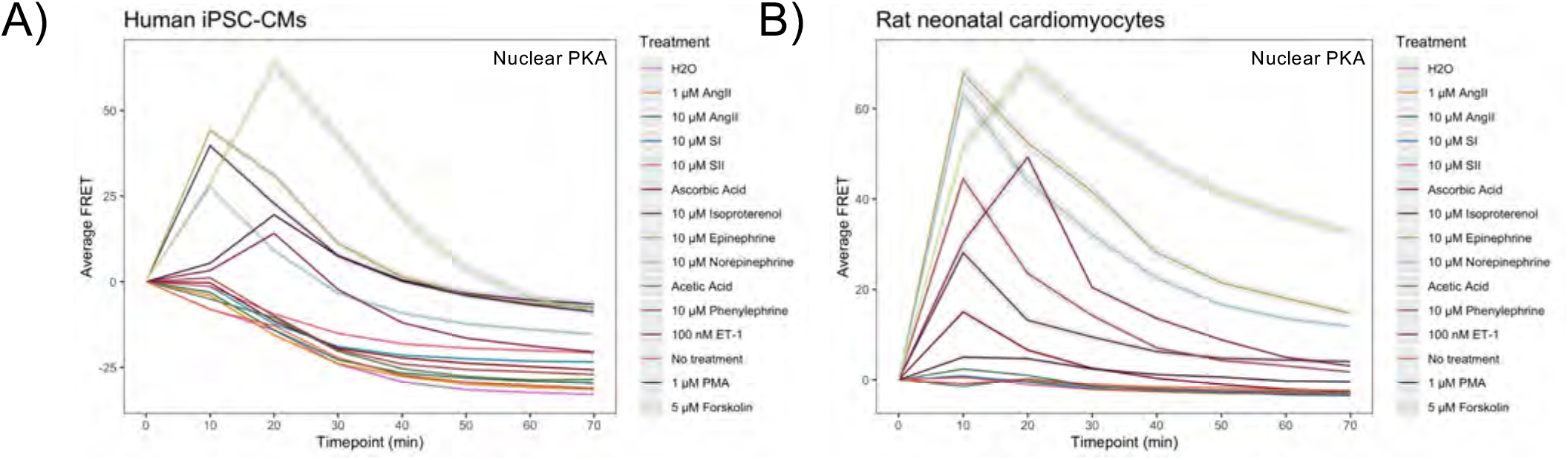
Mean nuclear PKA activity over a 70-minute timeframe. Baseline normalized averaged nuclear PKA responses measured in **(A)** human iPSC-derived cardiomyocytes (hiPSC-CMs) and **(B)** rat neonatal cardiomyocytes (RNCMs). Grey shadow represents SEM.

**Supplemental Figure 2.**
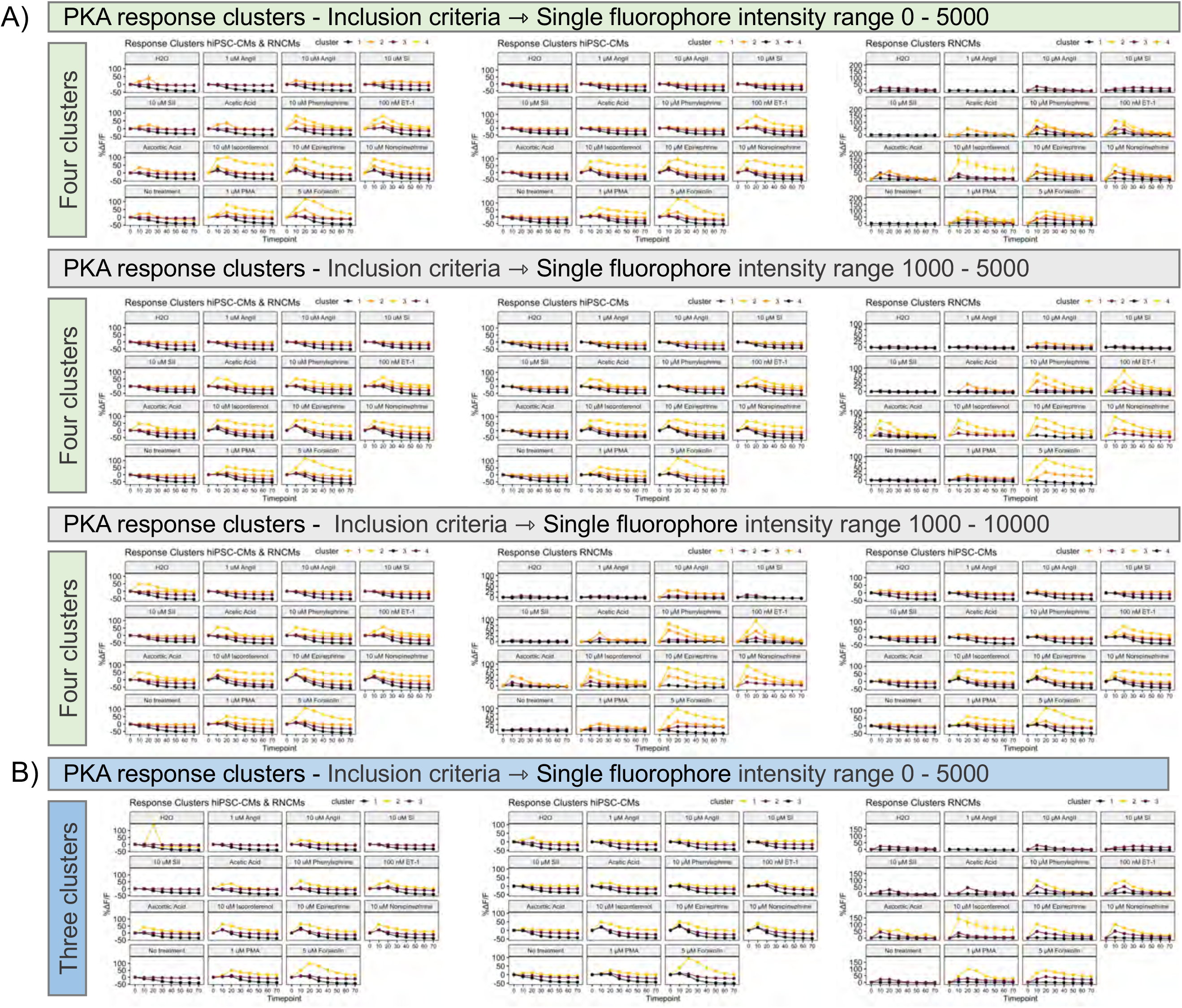
Graphical representations of PKA response clusters in hiPSC-CMs and RNCMs. Response clusters were plotted as a function of ExRai-AKAR2-NLS biosensor expression (RFU), ranging from **(A)** 0 – 5000, 1000 – 5000 and 1000 – 10 000. Biosensor expression did not appreciably influence the response clusters generated. Requesting the clustering algorithm to unearth four clusters instead of **(B)** three provided a better fit for the data.

**Supplemental Figure 3.**
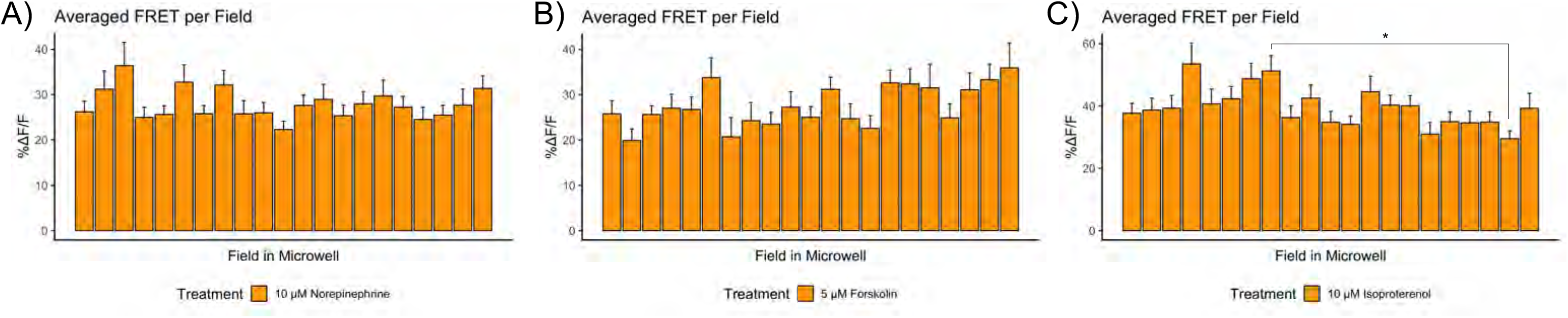
Bar plot representing drug diffusion patterns within different fields of imaged hiPSC-cardiomyocytes expressing ExRai-AKAR2-NLS. Charts display drug diffusion at the first timepoint post-stimulation with **(A)** norepinephrine (Welch’s ANOVA p = 0.3260), **(B)** forskolin (Welch’s ANOVA p = 0.0214), **(C)** isoproterenol (Welch’s ANOVA = 0.0043, Dunnett’s multiple comparisons test field 8 vs. 20 p = 0.0404), Error bars represent SEM.

**Supplemental Figure 4.**
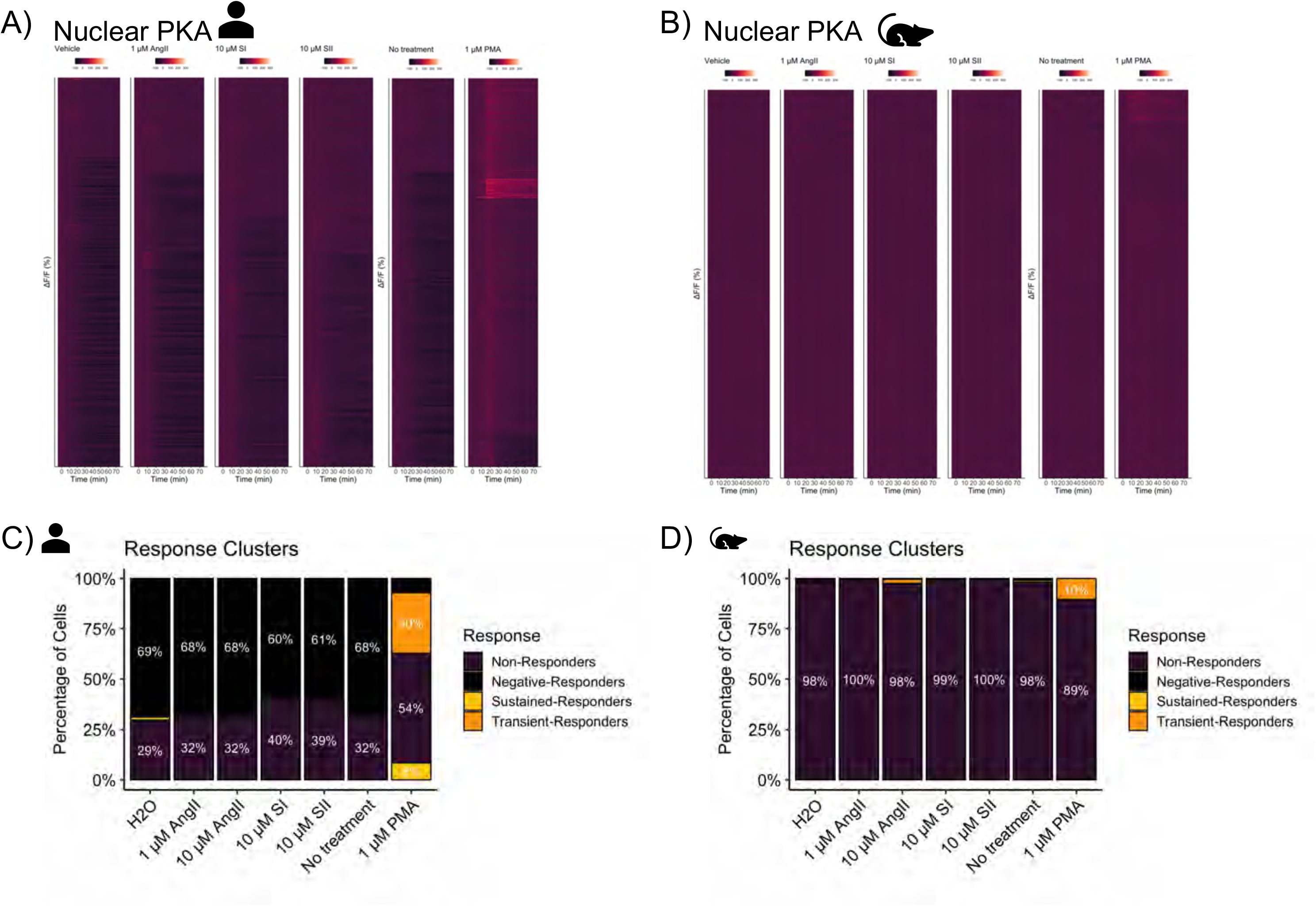
Single nuclei PKA activation patterns as observed in hiPSC-CM and RNCM cultures in response to angiotensinergic drugs. Heatmaps displaying nuclear PKA activity as measured in **(A)** hiPSC-CMs and **(B)** RNCMs. Single nuclear PKA data summarized as %ΔF/F (y-axis) as a function of time (x-axis). The data was partitioned into four clusters representing distinct nuclear behaviors, either exhibited sustained or transient responses to agonists while other nuclei failed to respond or experienced a decrease in activity compared to baseline. Representation of the four response clusters was plotted as a bar chart with percentages of nuclei belonging to each response cluster as observed in **(C)** hiPSC-CMs and **(D)** RNCMs. Experiments were performed using RNCMs isolated from 4 different neonatal rat pup litters and 3 independent cardiac differentiations.

**Supplemental Figure 5.**
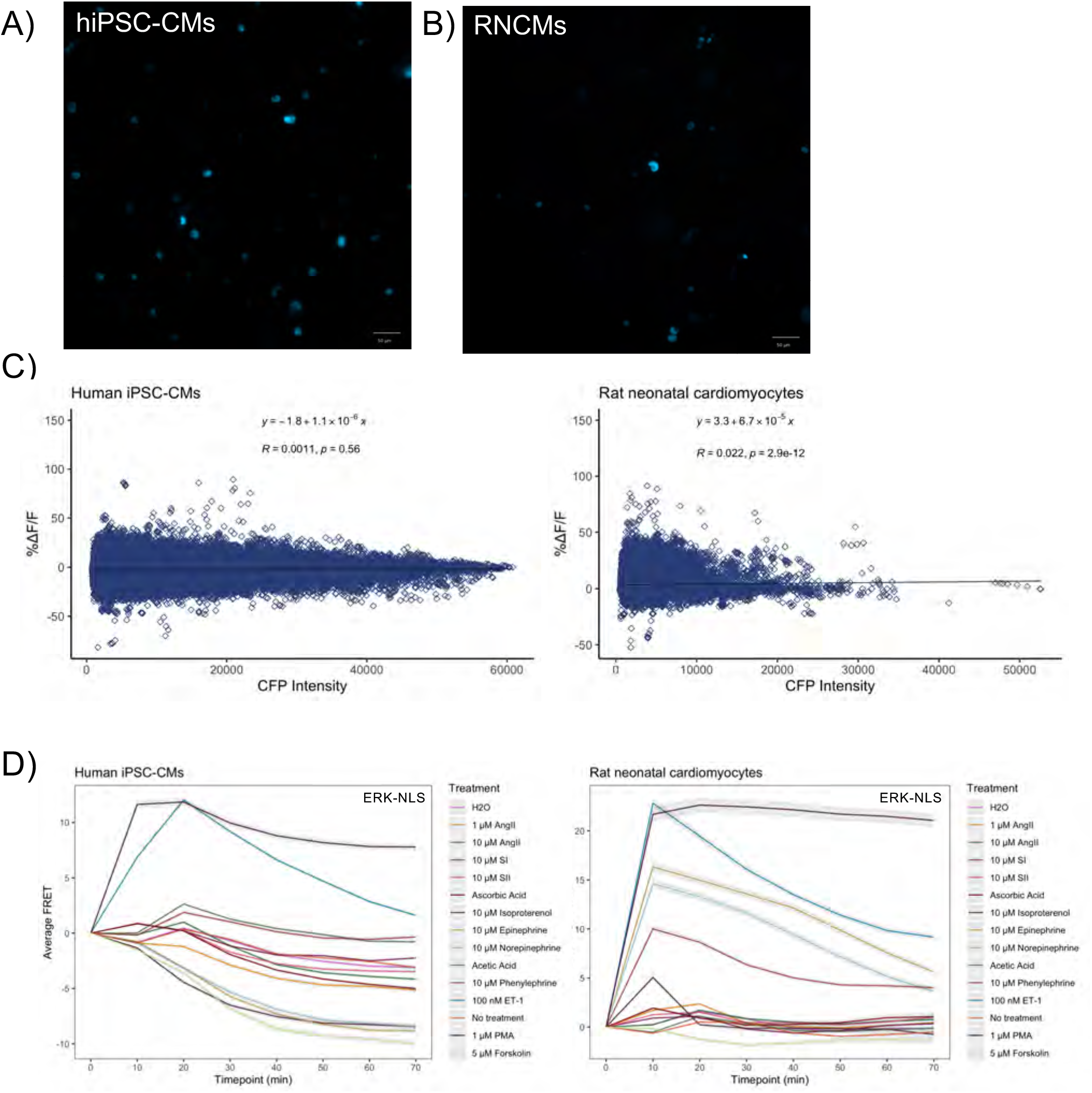
Nuclear ERK_1/2_ investigations in hiPSC-CM and RNCMs. Fluorescent microscopy image depicting expression of EKAREV-NLS donor in **(A)** hiPSC-CMs and **(B)** RNCMs. Agonist induced FRET responses plotted as a function of donor intensity in **(C)** hiPSC-CMs and RNCMs. Donor is expressed at higher intensities in hiPSC-CMs. **(D)** Averaged ERK_1/2_ response profiles in both cardiomyocytes cell types. Grey shadow represents SEM.

**Supplemental Figure 6.**
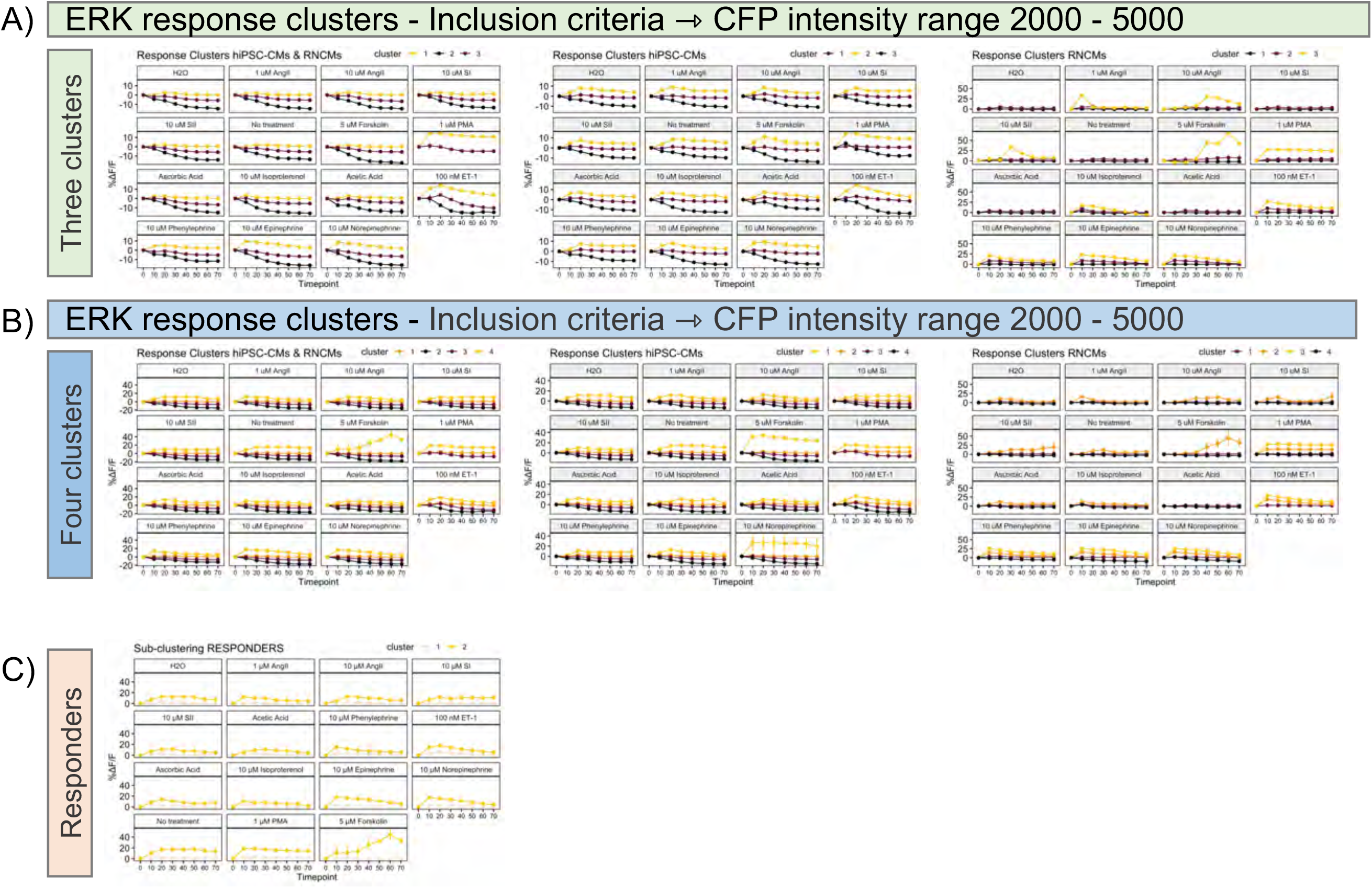
Graphical depiction of nuclear ERK_1/2_ response clusters. The clustering algorithm was requested to partition data in either (A) three or (B) four clusters. Based on the clustering output, nuclei appeared to experience three response profiles as opposed to four. This is exemplified in panel (B) as two clusters, displayed with orange and plum colours, appear to overlap with one another. **(C)** Depiction of the sub-clustering applied on the ‘Responder’ population. It is apparent that the ‘responding’ nuclei can be further divided into two clusters, referred as low- and high-responders.

**Supplemental Figure 7.**
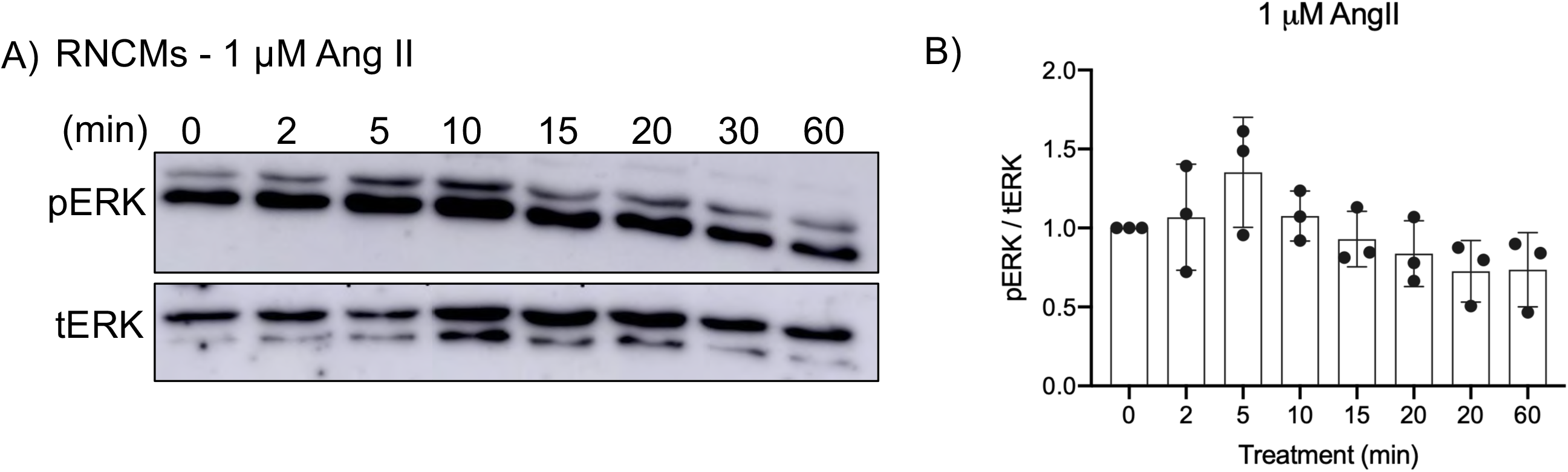
Ang II results in ERK_1/2_ action in RNCMs. Representative western blot demonstrating that Ang II stimulation results in ERK_1/2_ activity, depicted by pERK antibody at early timepoints, 5 minutes.

**Supplemental Figure 8.**
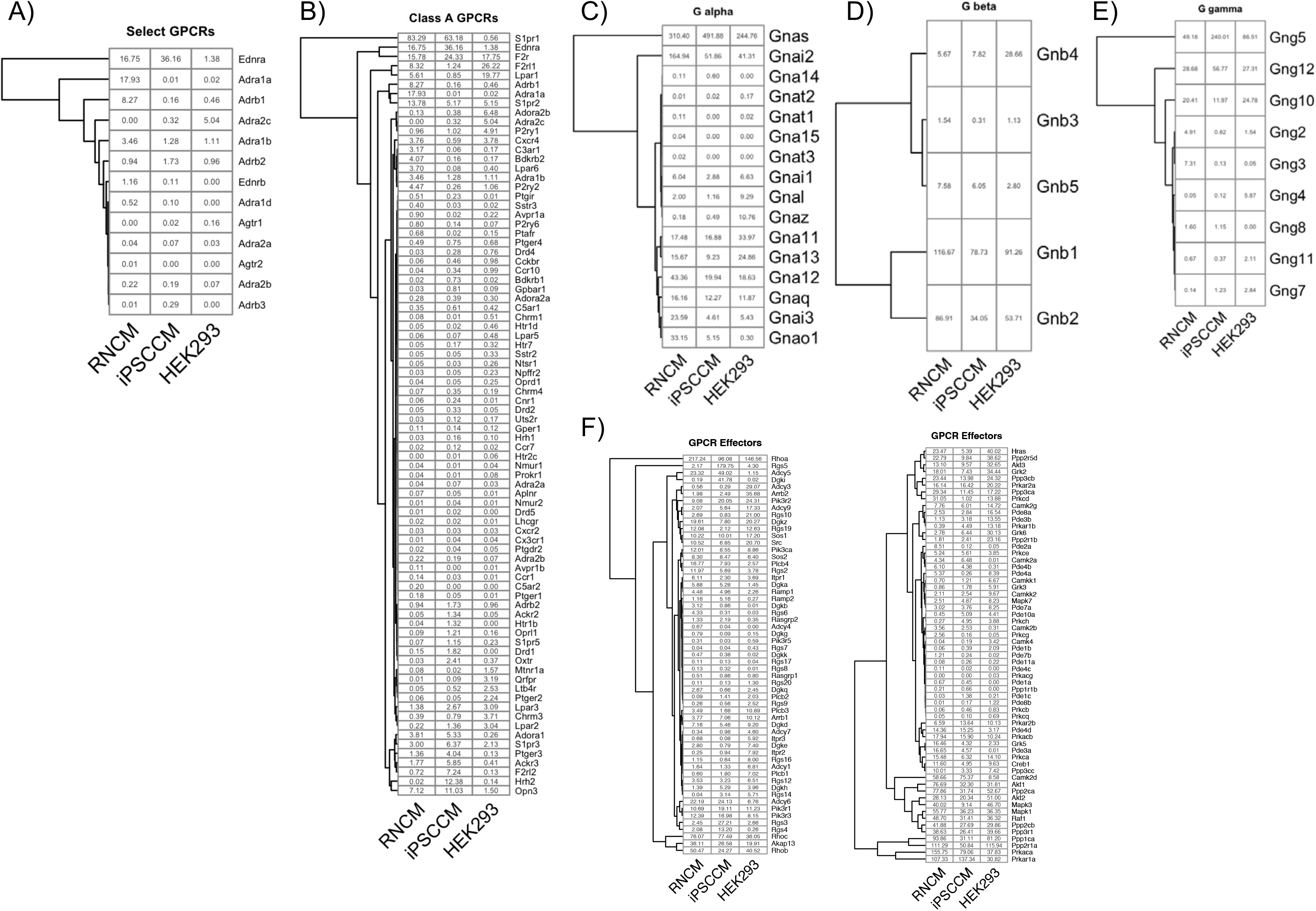
Ligand independent RNA-seq based comparison of the endogenous signaling machinery expressed in RNCMs, hiPSC-CMs and HEK 293 cells. Exploratory analysis of gene sets associated with GPCR signal transduction investigating **(A)** class A GPCRs, heterotrimeric G proteins **(B)** Gα, **(C)** Gβ and **(D)** Gγ as well as **(E)** effector expression profiles. Colorless heatmaps show transcript abundance measured in TPM, related to **Figure 5**. For comparison’s sake, data from HEK 293 cells was included (published originally as Lukasheva et al., 2020 Sci Rep 10, 8779. https://doi.org/10.1038/s41598-020-65636-3.

**Supplemental Figure 9.**
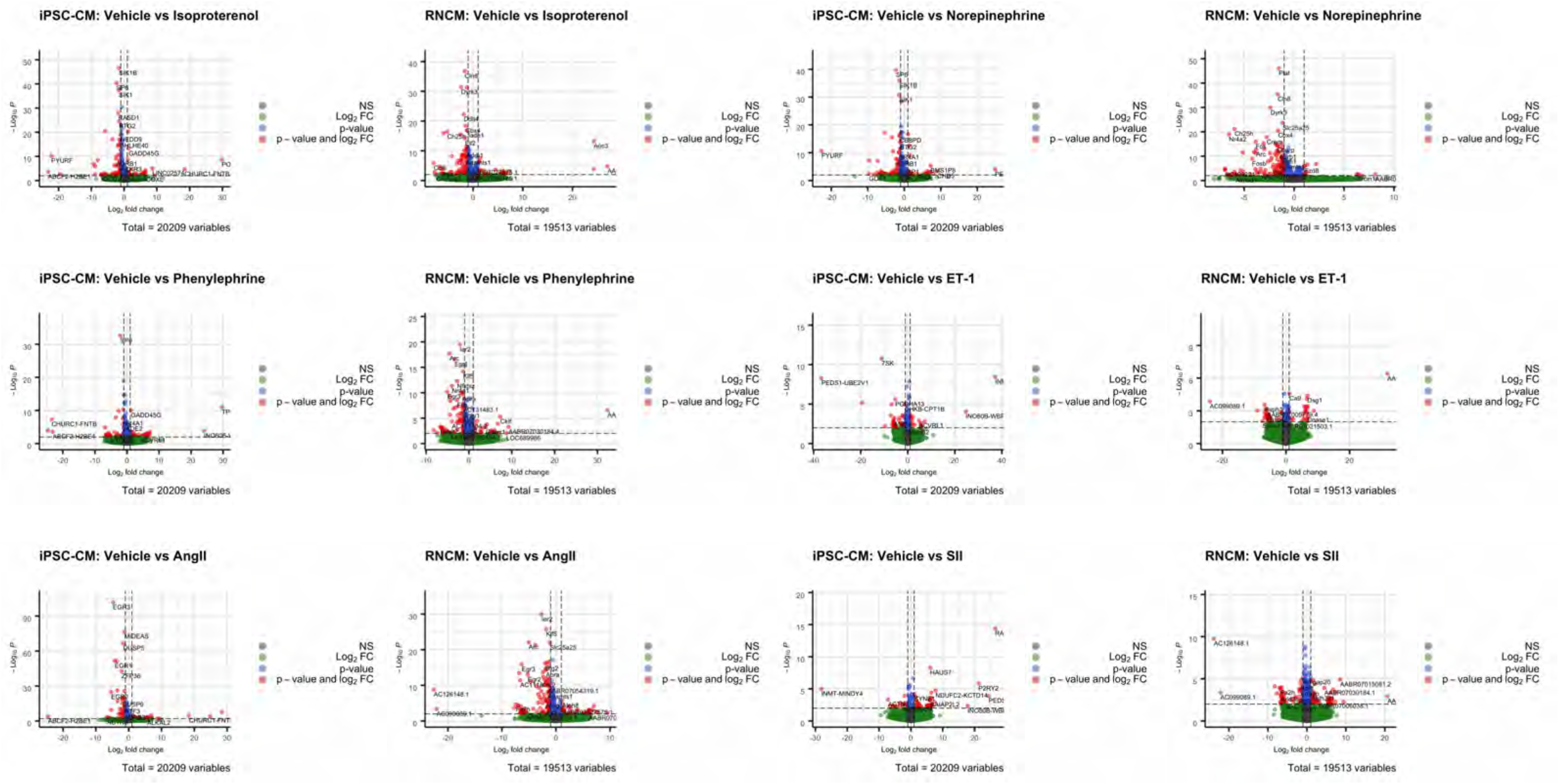
Volcano plots demonstrating up- and downregulated genes as determined by bulk RNAseq, 60 minutes after stimulation with various agonists.

**Supplemental Figure 10.**
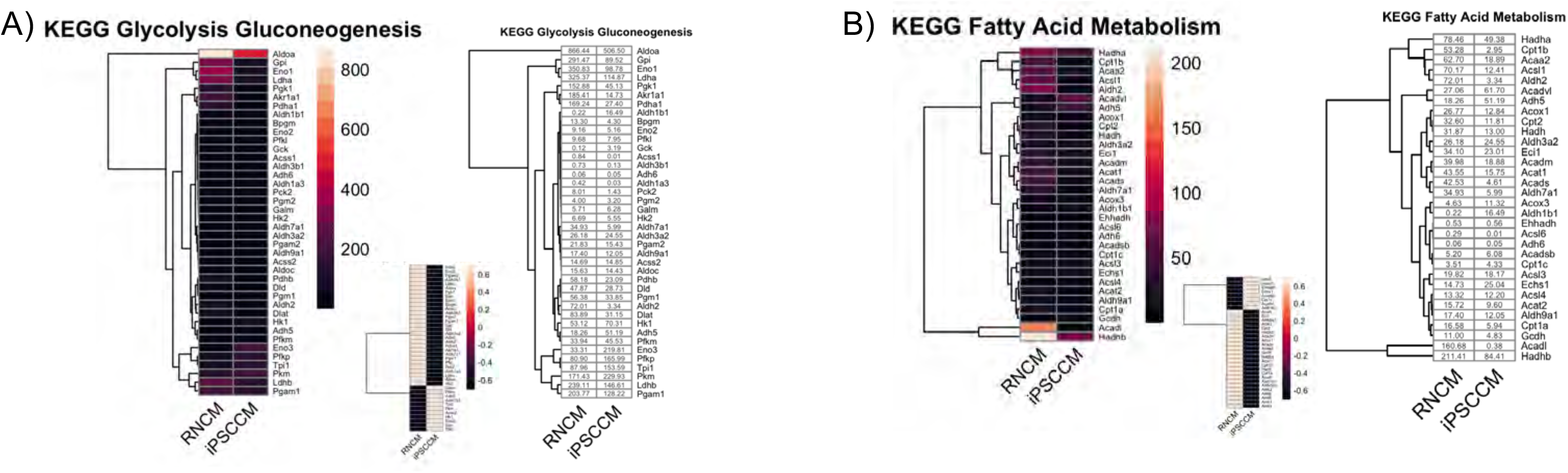
Transcript profiles of select energy metabolism-related gene sets. Transcript abundance (TPM) of gene involved in **(A)** glycolysis and **(B)** fatty acid metabolism. Large colored heat maps represent gene expression based on TPM while smaller colored heatmaps display comparative expression (hiPSC-CM vs RNCMs). Colorless heatmap depict raw transcript abundance.

## Acknowledgements

This work was supported by grants from the Canadian Institutes of Health Research (CIHR). K.B was supported by a doctoral studentship from CIHR and by a McGill Faculty of Medicine Internal studentship. J.J.T was supported by a doctoral studentship from the CIHR as well as the McGill Healthy Brains for Healthy Lives initiative. R.M. was supported by studentships from the McGill-CIHR Drug Development Training Program and the McGill Faculty of Medicine. We thank Dr. Michiyuki Matsuda (Kyoto University) and Dr. Jin Zhang (University of California) for providing us with biosensor constructs. We thank the McGill Advanced Bioimaging Facility (ABIF) for instrument access and user support. We also thank Nicolas Audet, the manager of McGill Pharmacology and Therapeutics Imaging and Molecular Biology platform for training and assistance with microscopy and image analysis. Likewise, we thank Jacob Blaney and Emma Paulus for their assistance in animal experiments. Lastly, we thank members of the Hébert lab for feedback and guidance throughout the development of the project, and for critical reading of the manuscript.

